# An agent-based model clarifies the importance of functional and developmental integration in shaping brain evolution

**DOI:** 10.1101/2020.05.04.075820

**Authors:** Shahar Avin, Adrian Currie, Stephen H. Montgomery

## Abstract

Comparisons of vertebrate brain structure suggest a conserved pattern of scaling between components, but also many examples of lineages diverging dramatically from these general trends. Two competing hypotheses of brain evolution seek to explain these patterns of variation by invoking either ‘external’ processes, such as selection driving phenotypic change, or ‘internal’ processes, like developmental coupling among brain regions. Efforts to reconcile these views remain deadlocked, in part due to empirical under-determination and the limitations of ‘relative significance’ debates. We introduce an agent-based model that allows us to simulate brain evolution in a ‘bare-bones’ system and examine the dependencies between variables that may shape brain evolution. Our simulations formalise verbal arguments and interpretations concerning the evolution of brain structure. We illustrate that ‘concerted’ patterns of brain evolution cannot alone be taken as evidence for developmental coupling, or constraint, despite these terms often being treated as synonymous in the literature. Both developmentally coupled and uncoupled brain architectures can represent adaptive mechanisms, depending on the distribution of selection across the brain, life history, and the relative costs of neural tissue. Our model also illustrates how the prevalence of mosaic and concerted patterns of evolution may fluctuate through time in a variable environment, which we argue implies that developmental coupling is unlikely to be a significant evolutionary constraint.

## Introduction

How are macro-evolutionary patterns in vertebrate brain structure best characterised, and what processes drive those patterns? Answering such questions requires understanding how developmental mechanisms or architectural constraints, as well as selection acting on neural traits, together shape and support behavioural and cognitive evolution. Debates over these conflicting pressures on variation have dominated vertebrate evolutionary neurobiology for decades, with no unified theoretical framework in sight.

At the heart of this debate are two views of brain evolution which, at their most polarized, make diametrically opposing predictions. Under one hypothesis, brain components are developmentally coupled such that the size of each component is largely determined by common developmental mechanisms, such as the timing and duration of neurogenesis (1–3). This would lead the entire brain to evolve in a ‘concerted’ manner, with the size of all components being closely predicted by overall brain size (1–4). Initially, this coupling was viewed as an evolutionary constraint (2) and, although proponents of this hypothesis now argue that developmental coupling is a mechanism that evolves in response to selection favouring conservative scaling (1,3), the view that concertedness in itself indicates developmental constraint remains widespread in the literature (e.g. 4–13). A contrasting hypothesis instead argues that variation in brain components is developmentally independent of both other brain structures, and of total brain size, allowing them to respond to targeted selection pressures in a ‘mosaic’ way (14–17). Mosaic evolution can lead to adaptive, non-allometric changes in brain structure, with stabilizing selection otherwise maintaining scaling relationships between co-evolving, functionally interdependent brain components (15,18). In essence, the mosaic model favours ‘external’ explanations that emphasize the role of selection in driving both independent phenotypic evolution *and* co-variation of brain structures, while concerted theorists favour ‘internal’ mechanisms which emphasise the role of developmental coupling as a route to maintaining scaling relationships (19).

Perhaps confusingly both hypotheses have at times been supported by analyses of the same volumetric data (e.g. 2,15,20). Proponents of the ‘concerted’ view of brain evolution point to consistent allometric scaling between brain components and total brain size as evidence of strong developmental integration across brain structures (1,2). Proponents of the ‘mosaic’ model instead point towards co-evolution between brain components that is independent of total brain size, and evidence for ‘grade shifts’ that indicate deviations in scaling between taxonomic groups, as evidence that brain components are caught between distinct selection pressures, and constraint from functional-interdependence (15). Distinctions between these hypotheses have become more nuanced, with the concerted hypothesis incorporating periodic restructuring of the brain (3,21). However, universally satisfactory tests of these hypotheses have proven elusive.

There are two key reasons for this deadlock. First, proponents of concerted and mosaic models are engaged in a ‘relative significance debate’ (22). Both sides agree that brain evolution exhibits features associated with both concerted and mosaic evolution, but disagree on which pattern dominates across evolutionary time, and why (see for example, 1, p. 299). Relative significance makes hypothesis testing difficult. Neither hypothesis is subject to critical tests, as both accommodate - and even expect - different degrees of departure from the ‘norm’. Competing views of brain evolution therefore run the risk of being too indeterminate for definitive testing.

Second, tests of these hypotheses are underdetermined by available evidence. Empirical support for concerted or mosaic evolution is most often drawn from comparative analyses of volumetric brain data. These data reflect the outcome of the interaction between competing adaptive and constraining factors, and do not, in themselves, provide evidence of the developmental mechanisms involved (23,24). This is a critical point, as ‘concertedness’ has frequently become a byword for developmental constraint (e.g. 4–13), biasing the interpretation and presentation of many studies. However, this is problematic because the mosaic brain hypothesis also predicts co-variation between interdependent brain regions. If the brain is viewed as a network of interdependent networks, these *functional* constraints could produce consistent scaling relationships across brain components— *i*.*e*. concerted evolution— without invoking developmental coupling (18). As inferred through classic evolutionary theory (25), merely recognising a concerted pattern is therefore insufficient evidence to assess competing mechanisms, or support the predominance of either hypothesis.

If patterns of phenotypic variation alone are unsuitable for identifying the mechanisms underpinning allometric scaling, what evidence could? As noted by previous authors, “it is not the *phenotypic* correlation that matters, so much as the *genetic* correlation” (26). Several recent quantitative genetics studies have found evidence of substantial genetic independence between brain components (5,7,27), a central prediction of the mosaic brain hypothesis (reviewed in 18). However, these studies typically concern standing genetic variation within populations. The developmental coupling hypothesis can accommodate this evidence if much of this genetic variation will be deleterious, and therefore does not reflect the genetic architecture favoured by selection over evolutionary timescales. Hence, neither phenotypic data nor currently available genetic data, are sufficient to unite views on the relative importance of developmental and functional coupling, constraint, and adaptive lability in the evolution of brain structure.

When faced with relative significance and empirical underdetermination, simple mathematical models can help realise basic causal dynamics in a ‘bare-bones’ system, and are a way of examining the dependencies between variables that are thought to be important. Such models can inform empirical studies by making explicit hitherto hidden assumptions, and revealing surprising emergent dynamics (28). This approach has recently been applied to debates over the socio-ecological selection pressures shaping brain size (29–32), providing a new approach to the field of evolutionary neurobiology. Here, we introduce an agent-based model of brain structure that allows us to explore the interactions between fitness and constraints derived from selection, development and function. In particular, our model allows us to formalise several verbal arguments over the role of developmental coupling in brain evolution:

1. Do functional dependencies produce concerted patterns of evolution as well as developmental coupling (c.f. 23,24)?
2. Do the costs of neural tissue select against concertedness when selection acts on a specific brain component (c.f. 33)?
3. Is developmental integration evolutionarily labile (c.f. 34), and do functional dependencies select for developmental coupling (c.f. 23,34)?
4. Does developmental coupling allow brains to evolve adaptively more quickly? (1)?

Our model allows us to explore these previously verbal arguments and interpretations of volumetric data. We demonstrate that this ‘bare-bones’ model helps clarify current debates surrounding the evolution of brain structure by capturing the basic evolutionary dynamics at play, and hope that it shifts these debates in a productive theoretical and empirical direction.

## Methods

To explore the interplay between developmental and functional coupling on the evolution of brain structure, we devised a model that simulates the evolution of a population of individuals in an environment, where fitness is determined by the size and costs of brain components.

### i) Definitions

We employ terminology from the evolutionary neurobiology literature, but the debate could also be understood in terms of principles derived from quantitative genetics. ‘Concerted evolution’ is the result of correlated evolution among all brain components, which can be caused by two main processes. The first, referred to as ‘developmental coupling’ occurs through genetic correlations that act through development, such that variation at a particular locus affects the development of multiple, or all, brain components. ‘Developmental de-coupling’ refers to the breakdown, or absence, of these genetic correlations. The second, referred to a ‘functional coupling’ instead occurs through correlated selection pressures, which either arise due to components contributing to shared behavioural or computational functions, or the environment selecting on multiple aspects of brain structure (e.g. dark environments selecting for increased olfactory processing and against visual processing; in this case the selection correlation is negative). Both correlated selection and genetic correlations can lead to correlated evolution (25) and our model is designed to explore this in the context of brain evolution. However, we note it is likely applicable to other anatomies.

### ii) Model components

The model is characterized by three components: the population of individuals, the environment, and an evolutionary algorithm. We provide a simplified description of the model in Figure S1.

Each individual is characterized by a number (*N*) of *brain components*, each component having a specific *size* (*S*_*i,j*_), where *i* denotes the individual and *j* denotes the *j*-th of *N* components. For example, among individuals with three brain components (*N*=3), the brain of ‘Individual A’ could have a first component of size *S*_*A,1*_=5, a second component of size *S*_*A,2*_=7, and a third component of size *S*_*A,3*_=2, which we denote as *S*_*A*_=(5, 7, 2). A second individual, ‘Individual B’, might have component brain sizes of 3, 3 and 1 respectively, denoted *S*_*B*_=(3, 3, 1). In this case the total size of Individual A’s brain would be 14, while the size of Individual B’s brain would be 7. Our model assumes that all brain components have the same fitness cost per unit size (see below), so it is total brain size that determines the evolutionary cost of an individual’s brain, while the size of individual components contribute differently to fitness benefits. Whether evolution is mosaic or concerted depends upon the factors influencing changes in brain component sizes over time. The sizes of brain components are allowed to vary, through mutation, and this variation is influenced by *developmental coupling* (*D*), which takes values between 0.0 (no developmental coupling, i.e. a fully ‘mosaic’ brain) and 1.0 (complete developmental coupling, i.e. a brain structure fully determined by total brain size). *D* is analogous to the strength of genetic correlations between components. For example, a *D* of 0.5, would indicate that 50% of the variation in each component is determined by variation in total brain size, and 50% of the variation in each brain component is independent of variation in both total brain size and other components. When a mutation event occurs, the program generates *N*+1 random mutation factors (*m*_*i*_), for example between 0.5 and for a 50% mutation step size, where there is one factor is for the whole brain (*m*_*0*_) and one for each component (*m*_*1*_ to *m*_*N*_), each brain component is then scaled by these mutation factors, with the variation in mutations affecting particular brain components being flattened depending on *D’s* value. For example, when *D* is 0, *m*_*0*_ for total brain size mutation will be multiplied by 0 but other mutation factors will vary independently according to a scaling factor (1 – *D*, i.e. unscaled when *D* is 0), whereas when *D* is 1 all mutation factors for individual brain components will be multiplied by 0 and only the mutation factor for total brain size will persist. Models with intermediate values of *D* fall between these extremes. This is determined according to the following formula.

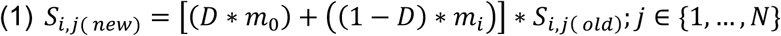

The *Environment* is characterized by three factors. First, a set of *benefits* (*B*_*j*_), one for each of the *N* brain components, which represents the fitness contribution of each size unit an individual gains from that particular brain component, at a particular size. For example, in an environment with benefits *B*=(1, 2, 3), ‘Individual A’ from above will have total fitness contributions from brain component sizes of 5*1 + 7*2 + 2*3 = 25, while ‘Individual B’ will have total fitness benefits of 3*1 + 3*2 + 1*3 = 12. Second, the environment imposes a fitness *cost* (*C*) per size unit of brain, which is uniform for all brain components based on the assumption that there is a linear ‘per neuron’ energetic cost (35), and that units of ‘size’ in the model is analogous to neuron number. Total fitness for an individual *i* with brain component sizes *S*_*i,j*_ in an environment defined by benefits *B*_*j*_ and cost *C* is thus given by ∑ _*j*_(*S*_*i,j*_ ∗ *B*_*j*_) - (*B* ∗ ∑ _*j*_ *S*_*i,j*_). For example, if the environment described above had a cost *C*=1 per unit component size, Individual A will experience a fitness cost of 14, giving a total fitness of 25 - 14 = 11, while Individual B will experience a fitness cost of 7, giving a total fitness of 5. Third, an environment is characterized by *Functional Coupling* (*F*), between 0.0 (no functional links between two brain components) and 1.0 (complete functional interdependence between two brain components), which determines how similar brain component benefits are to each other (for *F*=1, all benefits are identical).

The *Evolutionary Process* progresses through the following steps:

1. Determine the number of ‘offspring’ for each individual in a population, and the age of all individuals, measured in number of generations (these are identical across the population).
2. Initialise an environment and a population of individuals with identical, uniform brain component sizes (*S*_*i,j*_ = 1).
3. For a given number of simulation steps

a. Generate offspring for all individuals
b. Mutate all offspring
c. Increase the age of all individuals
d. Remove individuals whose age exceeds the maximum age
e. Rank individuals according to their total fitness in the environment (described above)
f. Remove low-rank individuals until the population size returns to the origin size (i.e. population size is stable over time).

Given the above model parameters, we can examine the effects of developmental coupling (*D*) and functional coupling (*F*) on the evolution of brain component sizes. We can also explore the evolution of brain structure in populations of individuals with an intermediate value of *D* (unless specified otherwise, we use *D=*0.5), which are neither fully-concerted nor fully-mosaic. We call these ‘partially-mosaic’ individuals, where some of the mutations affect total brain size, scaling each component equally, and some affect each component independently, as given by equation (1) above. We can then assess which mechanism, a fully mosaic brain (*D*=0), a fully concerted brain (*D*=1), or a partially-mosaic brain (*D*=0.5), is most successful in different scenarios by measuring the frequency of individuals in a population with that *D* value, as a proportion of total population size, after *n* generations. The model is currently implemented with no upper ceiling on overall brain size, it therefore most accurately simulates periods reflecting directional increases/decreases in brain components, and by extension brain size, which is a common but not universal trend (36,37). This means that under static selection regimes the model can lead to continuous directional changes in total brain size. However, our focus is on brain structure, not brain size *per se*, and additional effects of imposing an upper ceiling on brain size can be inferred from the results and are discussed below. We also allow the environment to change randomly over the course of a simulation to examine how temporal heterogeneity in selection regimes affects the long-term success of alternative brain models. The code files in which the model is implemented are available from github.com/shaharavin/BrainEvolutionSimulator. Biplots were made using PlotsOfData (38). Many simulations produce a bimodal distribution of frequencies, we therefore display the mean of these iterations solely to illustrate the skew in the outcome.

### iii) Model comparisons

We conducted a series of simulations to explore four key questions identified in the introduction:

1. What mechanisms can produce concerted evolution? Here we fixed *C* and *B* while varying *D*. We then ran the simulation to examine the probability of obtaining coordinated changes in brain components under high, moderate and low levels of *F*.
2. How do costs of brain components affect the way brains respond to selection? Here we fix *F* and *B*, and vary *C* under a range of conditions. Our aim was to test whether the costs of additional brain tissue alter the probability of obtaining a mosaic or concerted brain.
3. How does variation in fitness contributions from different components affect the way brains respond to selection? Here we fix *F* and *C*, and vary *B* for each brain component under a range of conditions. Our aim was to test whether different levels of variation the fitness contribution of additional brain tissue alter the probability of obtaining a mosaic or concerted brain.
4. In a fluctuating environment do strong developmental constraints evolve or collapse? In our final comparison we took a different approach. We introduced a starting population of brains with a range of component sizes and *D*. We then allowed *C* and *B* to vary randomly every 2 generations to test what combination of factors persist over time. We subsequently explored how varying key life history traits (numbers of offspring and maximum age) might buffer the effects of random environmental fluctuation

In each case, the simulation was run over 100 generations, with 1000 iterations, a fixed population size of 300 individuals, initiated with 100 individuals per *D* value. In the main text we present results of simulations with a mutation step size of 5%, but models were also run with a larger mutation step size of 50% to examine how effects were influenced by mutation size (full results are described in the Supplementary Information). To explore the early stages of the simulations, we also present results from a subset of simulations with 10 generations. In experiments 2 and 3 simulations were run with an initial population containing equal numbers of individuals with different *D* values (*D*=0, 0.5 or 1), which were then evolved under different environment conditions (determined by *F* and the ratio of benefits to cost). In experiment 4, environmental conditions were randomly varied every 2 generations for 150 generations. The ‘success’ of a *D-*value was determined by the proportion of individuals in a population with that *D* value at the end of the simulation run. The full output of all models are summarised in Figures S2-S19. The frequency of *D* values were compared using generalised linear models and the glm() function in R (39) across batches of nine, 1000-iteration simulations where *F, D* and *B and/or C* varied (experiments 1-3, see for example Figures S2,S6), or where *F, D*, maximum lifespan and offspring number varied (experiment 4, see, for example Figures S14) (all experiments total *n=*27,000 iterations). We estimated the effects of all parameters and interactions where indicated, with a Gaussian distribution when comparing ‘degrees of mosaicism’ (defined as the natural log of the ratio between the largest brain component and the smallest brain component in each individual, averaged across the population) and a quasibinomial distribution when comparing proportional frequencies.

## Results

### i) Developmental and functional coupling can both produce concerted evolution

By varying the degree of both developmental coupling (*D*), and functional interdependence (*F*), between brain components, our model suggests that patterns of ‘concerted’ brain evolution, in which the size of brain components are correlated, can be caused by both developmental and functional coupling (Figure 1, Figure S2-4). Unsurprisingly, the probability of patterns of concerted evolution declines with decreasing *D* (*t*=50.330, p<0.001; Figure 1D). However, there is a significant interaction between *F* and *D* (*t*=28.730, p<0.001) whereby, for a given value of *D*, where *D* < 1, the probability of concerted evolution increases with higher values of *F* (Figure 1A-C, D), demonstrating that functional interdependence also promotes concertedness. Even when components are completely developmentally independent (*D* is set to 0), high values of *F* can favour comparatively low levels of mosaicism (Figure 1C). Mutation size also has an effect on the outcome, with more mosaic brains evolving with larger mutation step sizes for a given *D* value (*t*=133.650, p<0.001; compare Figures S2 and S3). These results illustrate that macro-evolutionary patterns of allometric scaling consistent with concerted evolution are not, in themselves, sufficient to distinguish between alternative mechanistic models of brain evolution.

**Figure 1:**
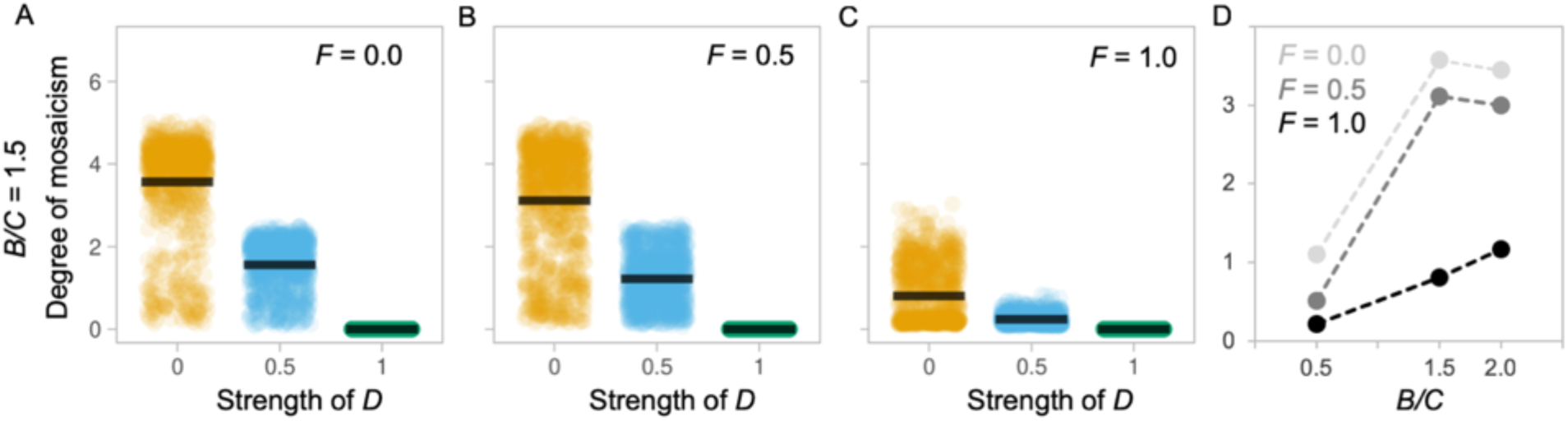
Evolution of ‘mosaicism’ under alternative conditions. **(A-C)** Each plot depicts the ‘degree of mosaicism” (y-axis, defined as the natural log of the ratio between the largest brain component and the smallest brain component in each individual, averaged across the population) as a function of developmental coupling *D* (x-axis) at the end of a 100-generation simulation run, compiled over 1,000 simulation runs, for a population of 100 individuals with an identical *D*, under different environments, defined by their functional coupling *F*, with a benefit to cost ratio (*B/C*) of 1.5 and a mutation step size of 5%. **(D)** Summary of the effects of varying *F* and *B/C* on the degree of mosaicism. See Figures S2-S4 for full results varying *B/C* and *F*, iteration numbers and mutation step size.

### ii) Mosaic and concerted evolution can both be adaptive

By competing alternative *D* values against one another, we found that both *D* (*t*=21.358, p<0.001) and *F* (*t*=42.595, p<0.001) affect survival in a particular environment. At an intermediate benefit to cost ratio (*B/C*=1.5) we found that the probability of success of a mosaic, low *D* value population increases when *F* is low, while high values of *F* result in a higher probability of success for the concerted, high *D* value population (Figure 2A-C; Figure S6-S8). A ‘partially-mosaic’ population (*D=*0.5) is very rarely favoured (Figure 2A-C; Figure S6-S8). These comparisons indicate that variation in selection across components alters the outcome of competition between populations with different levels of developmental coupling.

**Figure 2:**
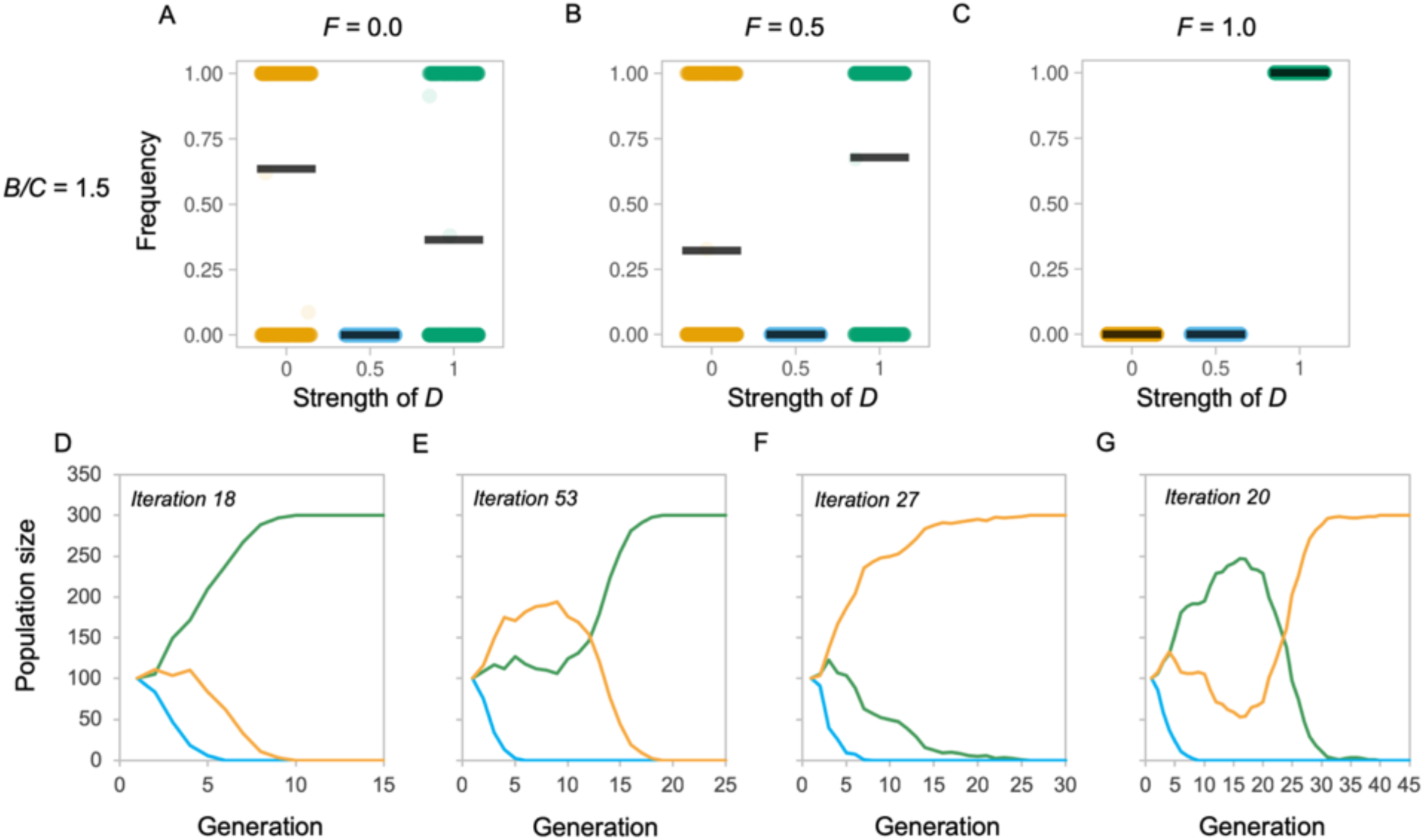
Selected examples of competition between evolving populations with different *D* values. (A-C) Each plot depicts the frequency of that *D* value relative to the total population (y-axis) as a function of developmental coupling *D* (x-axis) at the end of a 100-generation simulation run, compiled over 1,000 simulation runs, under alternative environments defined by their functional coupling *F*, with a benefit to cost ratio (*B/C*) of 1.5 and a mutation step size of 5%. Populations are initialised such that there are 100 fully mosaic individuals (D=0.0), 100 partially-mosaic individuals (*D*=0.5) and 100 concerted individuals (*D*=1.0). (D-G) Selected, representative, individual simulations showing the change in population frequencies over generations for a 5% mutation size and *B/C*=1.5. Colours indicate the *D* value, where yellow is *D*=0, where blue is *D*=0.5, and where green is *D*=0. These show a general pattern of smooth progression from the starting state of equal populations to one *D* value winning out (D, F), with only a minority of iterations showing signs of populations ‘swapping’ the lead (E,G). This consistency is expected under constant selection regimes. See Figures S6-S8 for full results varying *B/C* and *F*, iteration numbers and mutation step size, and Figures S9-11 for simulations in tailored environments that illustrate parameter effects.

### iii) Costs of additional tissue alter the outcome of competing models

In experiment 1, the degree of mosaicism is also associated with *B/C* ratio (*t*=131.790, p<0.001; Figure 1D, Figure S1), which interacts with *D* (*t*=-90.970, p<0.001) such that the degree of mosaicism tends to increase with *B/C* (Figure 1D). We next repeated the simulations described in (ii) while varying the *B/C* ratio associated with additional brain tissue. The initial models were run with an average *B/C* ratio of 1.5 (Figure 1, Figure S6-8 second row), and were re-run with ratios of 2 and 0.5 (Figure 3A-C, S6-8, first and third rows), simulating low and high costs of brain tissue. This revealed tissue costs have a major effect on the success of populations with different *D* values (*t*=12.116, p<0.001). *B/C* interacts with *D* (*t*=-13.885, p<0.001) such that, for a given combination of *F* and *D*, a high *B/C* (=2) consistently increases the probability of success for the population with a high *D* value (Figure 3A-C, Figures S6-S8). In contrast, for a given combination of *F* and *D*, a moderate *B/C* (=1.5) consistently increases the probability of success for low D, mosaic models. However, when the *B/C* ratio is low (=0.5) the non-linear nature of the interaction between *C, F* and *D* is revealed, such that concertedness again becomes successful, even at low *F* values (Figure 3D-F, Figure S6-S8 third rows). In these competition experiments, mutation size had no effect on the outcome (*t*=0.740, p=0.459; compare Figures S6 and S7). The non-linear success of low *D* values can be explained if developmental coupling facilitates rapid *decreases* in all brain regions when costs of brain tissue exceed the benefit, allowing a quick escape from a costly phenotype, or when the fitness of the whole system is dominated by a single component, such that increases in total brain size approximate the fitness benefit of increasing specific components.

**Figure 3:**
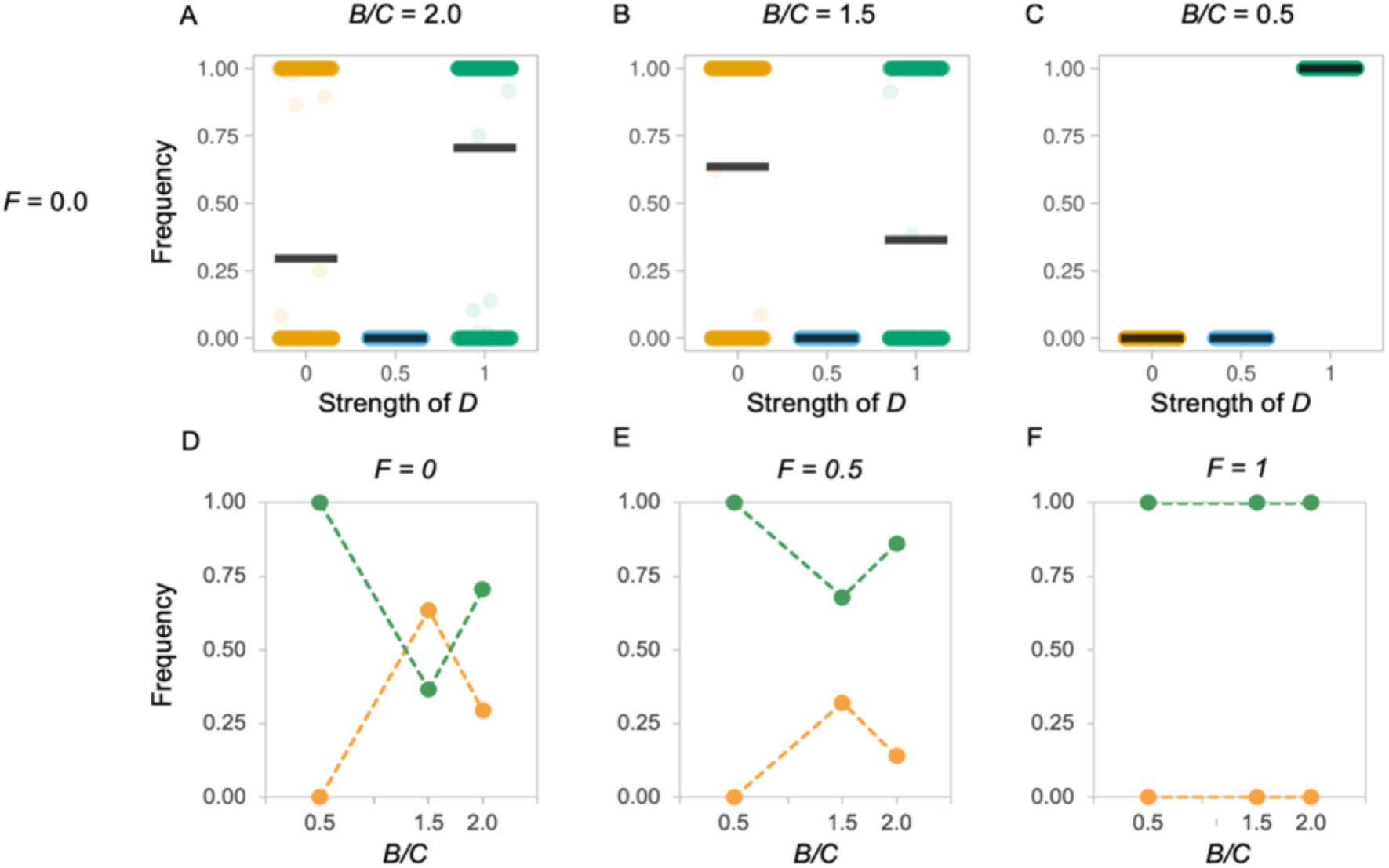
Selected examples of competition between evolving populations with different *D* values, showing the effects of varying the benefit to cost ratio (*B/C*). **(A-C)** Each plot depicts the frequency of that *D* value relative to the total population (y-axis) as a function of developmental coupling *D* (x-axis) at the end of a 100-generation simulation run, compiled over 1,000 simulation runs, with a mutation step size of 5%, and *F*=0 to exemplify the effects of varying the *B/C* from 0 to 2. Populations are initialised such that there are 100 fully mosaic individuals (*D*=0.0), 100 partially-mosaic individuals (*D*=0.5) and 100 concerted individuals (*D*=1.0). (**D-E**) Each plot depicts the average relative frequencies of *D=0* (yellow) and *D=1* (green) at the end of a 1,000 iterations of a 100 generation simulation, at three *B/C* ratios, across three *F* values representing low (**D**), moderate (**E**), and high (**F**) functional coupling. See Figures S6-S8 for full results varying *B/C* and *F*, iteration numbers and mutation step size, and Figures S9-11 for simulations in tailored environments that illustrate parameter effects.

To further illustrate these effects, we specified particular parameter comparisons that show how *B/C* interacts with changes in selection regimes such that the most successful *D* value switches based on changing the benefit cost ratio (Figure 4, Figures S10-S12 rows) or variation of selection across components (Figure 4, Figure S10-S12 columns). In particular, we note that i) populations can maintain multiple populations with different *D* values where fitness is dominated by the contribution of one component (Figure 4A); ii) shifts in the probability of a population of a mosaic, low *D* population being successful are otherwise associated with increased variation in the *B* value among components (Figure 4Bii); and iii) the relative success of a mosaic, low *D* population increases when one or two components provide a net benefit to size, while other component(s) provide a net cost (Figure 4Biii), while populations with high *D* values are favoured when all components provide *either* a net benefit or a net cost (Figure 4Bi).

**Figure 4:**
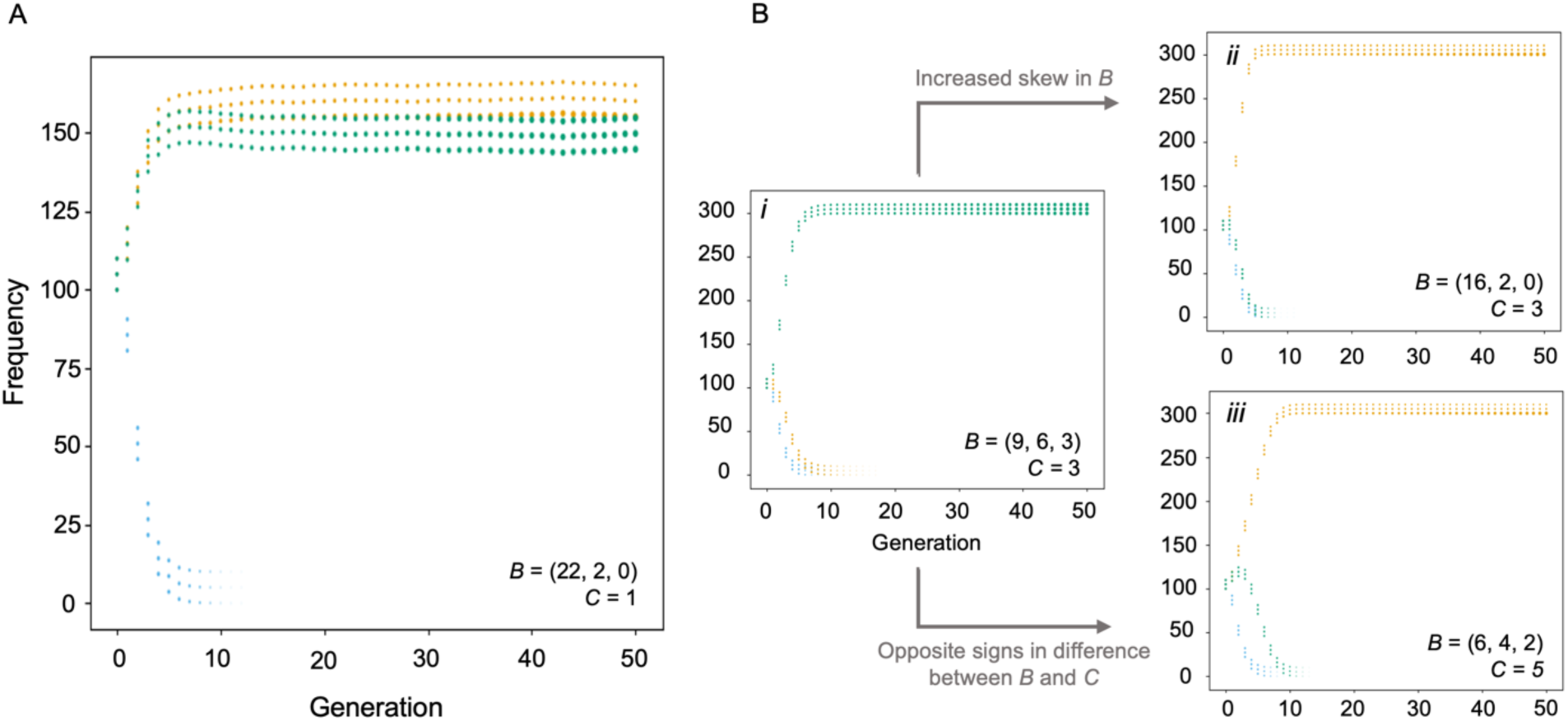
Selected examples of competition between evolving populations with different *D* values, showing the average frequency of each population during the first 50 generations of 1000 simulated runs, with “hand-crafted” environments depicted in the figure to illustrate specific situations of interest. Each plot depicts the average relative size of the 3 components within the artificial brain, with *D* values colour coded (*D*=0 in gold, *D*=0.5 in blue, *D*=1.0 in green). (**A**) When the benefits of each components are very strongly skewed, both *D*=0 and *D*=1 values can persist as each rapidly adjusts to approximate the optimal condition. (**B**) With moderate levels of variation in *B* across components a *D*=1 value spreads in the population (i), with increased skew in *B* across components the probability of spreading switches to *D*=0 (ii) unless skew is extreme in which case *D=1* again becomes successful (A). Shifts in the probability of success for *D*=1 are also associated with components having opposite signs in the *B/C* difference across components (ii).

### iv) Variation in the distribution of selection across time alters the outcome of competing D values, and interacts with life history

In a final set of simulations, we examined whether populations with alternative *D* values persist when the selection regime is temporally variable (due to randomised, independent changes in *B, C* and *F*). Under initial conditions, where offspring number was set to 1 and maximum age was set to 3, both concerted and mosaic populations persist over 150 generations with roughly equal probabilities (Figure 5A). Plotting the frequency of *D* values from individual simulations shows that the success of each population can fluctuate over time (Figure 5E-G), with multiple populations persisting, on average, for 62 generations (Figure S13A). This is substantially more than is found in simulations with fixed environments (Figure S5), and is also reflected in the high average generation at extinction for each *D value* (*D=*0, 91; *D=*0.5, 57; *D=*1, 98; Figure S13). We subsequently varied maximum age and offspring number to explore how ‘slow’ (long lives, few offspring) or ‘fast’ (short lives, many offspring) life histories buffer the effects of environmental heterogeneity. This revealed that the main effect captured in the model was that of offspring number interacting with *D* (*t*=43,622, p<0.001), with higher offspring numbers increasing the probability of success of the concerted, high *D* value (Figure 5A,C,D; Figure S14-S16 columns). Increasing maximum lifespan had a significant but smaller effect in the opposite direction (*t*= -4.349, p<0.001; Figure 5A,B,D; Figure S14-S16 rows). Altering the amplitude of fluctuations in the environment also has a subtle effect on the probability of success of competing populations with different *D* values (*t*=-8.986, p<0.001), with more extreme conditions slightly increasing the success of concertedness (Figure S18). In these competition experiments, mutation size again had no effect on the outcome (*t*=1.000, p=1.000; compare Figures S14 and S15).

**Figure 5:**
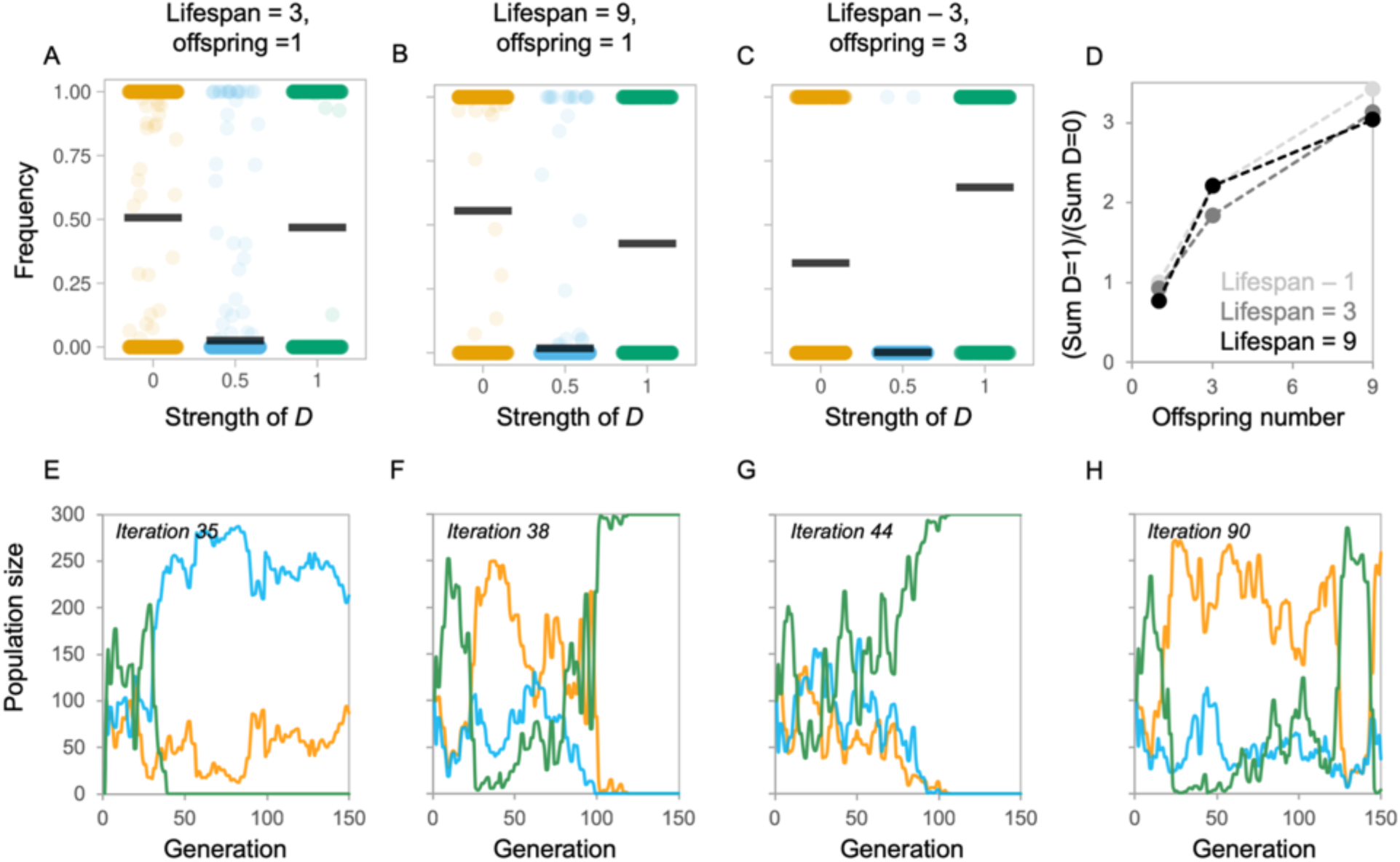
Selected examples of competition between populations with different *D* values in a randomly varying environment. **(A-C)** Each plot depicts the ratio of each population to the total population (y-axis) as a function of developmental coupling *D* (x-axis) at the end of a 150-generations, compiled over 1,000 simulation iterations; every 2 generations the environment was replaced, using a uniform random distribution for cost [0.5, 5], max average benefit [0.51, 105], and *F* functional coupling [0, 1]. Populations are initialized such that there are 100 fully mosaic individuals (*D*=0.0), 100 partially-mosaic individuals (*D*=0.5) and 100 concerted individuals (*D*=1.0). Simulations were performed varying two life history conditions: maximum lifespan, or the number of generations an individual persists alive, and offspring number, which are produced once every generation. Three combinations of lifespan and offspring are shown, for full comparisons see S13. **(D)** A summary of the ratio of concerted individuals (*D*=1) to mosaic individuals (*D=*0) at the end of each iteration of the 150 generation simulation, showing the effects of maximum lifespan and offspring number. **(E-G)** Selected, representative, individual simulations showing fluctuations in *D* value frequencies over generations. Colours indicate the *D* value, or *D* value, where yellow is *D* =0, where blue is *D*=0.5, and where green is *D*=1. See Figures S13-S16 for full results varying *B/C* and *F*, iteration numbers and mutation step size, and S17 for effects of increasing the size of environmental fluctuations.

## Discussion

The results of our agent-based simulations have several implications for studies using comparative volumetric data to interrogate mechanistic hypotheses of how brains evolve. First, they demonstrate that patterns of ‘concerted’ evolution (consistent allometric scaling of all brain components across macroevolutionary time) should not be taken as evidence for developmental coupling, as concertedness can also evolve due to high functional interdependence in the absence of developmental coupling. Second, the most probable route to adaptive brain evolution is strongly influenced by whether the change in selection regime is skewed towards one brain component or is evenly distributed across the whole brain, and what the relative costs and benefits of increased investment in brain tissue are. Third, depending on the context, both concerted and mosaic evolution can be adaptive. Finally, our model shows that variation in selection regimes can result in both mechanisms being maintained in a population. Given these results, pluralism is a tempting settlement. However, this ecumenism opens new questions: Under what conditions do developmental coupling and un-coupling succeed? What causes switches between mosaic and concerted modes of evolution? And how can we empirically distinguish them?

### i) Concertedness is a phenotypic pattern devoid of mechanistic information

Our simulations clearly show that concerted brain evolution can occur under any level of developmental coupling, or *D* value (Figure 1). This is consistent with classic quantitative genetics frameworks, which show that correlated evolution does not depend on strong genetic correlations among traits (25). However, despite early arguments that this is the case (23,24,26), the distinction between concerted, or correlated, evolution and developmental constraints is often neglected in the debates surrounding the evolution of brain structure. In our model, the probability of concertedness predictably increases with *D*, but it also increases with *F*, even when *D* is zero. Why do high *F* values result in concerted evolution? If we assume that selection acts on specific brain components, and excess brain tissue is generally expensive, then selection to maintain functional relationships should be expected, and would result in co-evolution among brain components. These formal results are consistent with empirical data. For example, across mammals the major components of the visual processing pathway, including the peripheral visual system, lateral geniculate nucleus and visual cortex, tend to co-evolve with one another, as predicted by their functional integration (40–42). Similarly, major components of the olfactory pathway, including the olfactory bulbs and olfactory cortex, also co-evolve (41,43). However, the visual and olfactory pathways show no consistent pattern of co-evolution between them (41).

Indeed, whether they co-vary negatively, positively, or not at all, can be explained by how diet and activity pattern interact to shape foraging behaviour (41). In addition, major brain structures connected by long-range axons, such as the neocortex and cerebellum also tend to co-evolve independently of total brain size, while also showing evidence of temporally transient independent change (44–47). These examples illustrate the effects of functional integration on co-evolution among brain components. Finally, our model demonstrates an interaction between *D* and *F*, which may suggest that a pattern of concertedness driven by *F* could select for, or against, developmental integration. Regardless, our results demonstrate that concerted patterns of brain evolution provide no evidence either for or against the prevalence of developmental coupling.

### ii) The costs and benefits of brain tissue, and the complexity of the mutation landscape

Our simulations suggest that the cost of excess brain tissue and the distribution of selection across networks of brain components play critical roles in determining how brains evolve. When the strength and direction of selection is skewed towards one brain component, the probability that a population with mosaic brains, or low *D* values, will be successful is increased. However, when the relative costs of neural tissue are either very low or high, the balance can switch to favour a high *D* values. This likely reflects the ‘speed’ at which the two populations can respond to selection. With only one mutational mechanism, it is potentially more likely for a developmentally coupled brain to evolve towards a new ‘adaptive peak’ than it is for a mosaic brain. This is because the mutational landscape for a mosaic brain is more complex, and the probability of hitting the optimal mutational path is reduced by a greater number of potential mutational outcomes.

At face value, our simulations therefore support previous arguments that “…a coordinated enlargement of many independent components of one functional system without enlargement of the rest of the brain may be more difficult, as its probability would be the vanishingly small product of the probability of each component enlarged individually” (2, p. 1583). However, if the costs of brain components are unequal, as may be expected if energetic consumption scales with neuron number (35) and neuron density varies between components (48), then this effect would be dampened according to the distribution of costs and benefits across the whole system. Hence, an uneven distribution of neural costs would likely increase the probability of mosaicism. Evidence that this effect occurs in nature may be provided by recent data showing life history traits constrain specific brain regions independently of overall brain size (49). In addition, our model currently imposes no upper limit on total brain size. While this will reliably reflect periods of directional changes in brain, or component, size (36,37), the close correlation between brain and body size, which may evolve under contrasting selection pressures (25), may impose limitations on how brains respond to selection that are not captured in the current model. However, this can be envisaged in terms of non-linear relationships between the size of brain components and their costs and benefits. As brain size approaches an upper ceiling the cost of increasing each component is likely increased, resulting in increased disparity of *B/C* ratios for each component when selection is favouring increases in particular brain regions. As such, inclusion of an upper limit to brain size would increase the range of conditions under which a mosaic pattern of evolution is adaptive.

While accurately assessing the fitness costs and benefits of brain tissue remains challenging, we suggest some tentative predictions based on these interpretations. First, we expect mosaicism to be less likely during transitions to high quality diets, as the cost of neural tissue relative to the overall energy budget may be reduced. Second, under sudden periods of energy resource limitation brains should shrink, at least initially, in a concerted manner. Third, mosaicism may be more likely when energy intake is relatively constant, but selection favours changes in neural investment which involve energetic trade-offs. This may be common during periods of ecological change that are not associated with changes in body size (and by proxy energetic intake), or in taxa that are size limited.

### iii) Partially-mosaic brains and the maintenance of diversity

It may be surprising that partially-mosaic individuals, which are seemingly a compromise-state, generally have low probability of success in our simulations. This can again be explained by the ‘cost’ of mutation. In short, partially-mosaic individuals incur the highest cost of mutational complexity. If the direction of mutation is random, in a given generation the partially-mosaic brain simply has lower odds of increasing its fitness because mutations affecting whole brain and region-specific size may often be in conflict. The relative lack of genetic variation contributing to both whole brain and region specific size in natural populations (5,7,27) is in keeping with this conclusion. However, how mutational effects interact during brain development requires further investigation.

However, our simulations across temporally variable selection regimes indicate that populations with alternative *D* values can co-persist within a single population for many generations. This provides an alternative route to mixed temporal patterns of concerted and mosaic evolution within a single population. If we view *D* values in our model as alternative genotypes, this implies that genetic variation affecting specific components or total brain size may both either persist at low levels in natural populations or periodically arise *de novo* and spread through a population. Regardless, this would enable selection to fluctuate between favouring mosaic and concerted mechanisms, permitting both adaptive restructuring and adaptive conservation of brain structure.

### iv) Reconciling mosaic and concerted views

Our simulation results suggest a way of reconciling mosaic and concerted views of brain evolution. Developmentally coupled brains evolve in scenarios involving some combination of tissue costs being evenly distributed, and an extreme and variable fitness landscape, while mosaic brains are the result of environmental stability coupled with differentiated selection among components. The quick evolutionary response enabled by developmentally coupled brain evolution makes it ideal for circumstances where getting it ‘approximately correct’ quickly is advantageous, while mosaic evolution is favoured when accuracy trumps speed. This is also tied to life-history and environmental heterogeneity. For example, large offspring number, which increases short term competition, favours the fast response provided by developmental coupling, while lower competition allows mosaic populations to persist, find the optimum brain structure, and out-compete developmentally coupled individuals.

The contrasting benefits of concerted and mosaic evolution brings us back to a major initial division between the two hypotheses (1–4), which continues to be widely reported in the literature (e.g. 4–13); where developmental coupling does occur, is it a constraint, or has it evolved and is therefore *evolvable*? Our simulations support developmental coupling as a scenario-dependent adaptive mechanism, rather than a constraint. First, our simulations show that the fate of the two opposing mechanisms can vary through time. In nature, this would be reflected in the formation and breaking of genetic correlations between traits. Second, the general absence of genetic correlations observed in quantitative genetics studies is reconciled with concertedness via developmental coupling by invoking the importance of *de novo* mutation, but our simulations suggest that the probability that these mutations will spread to fixation depends on the environmental context. By showing that the distribution and size of costs and benefits are critical in determining the outcome of competing mechanisms, our model supports a view whereby global selection regimes determine the constraints acting on brain evolution, not the developmental program. Two key conclusions from this work are therefore i) patterns of concertedness should not be equated with developmental coupling, and ii) developmental coupling should not be equated to developmental constraint.

### v) Conclusions

In sum, our agent-based simulations of competing views of brain evolution provide a number of informative predictions that should refresh our view of this ongoing debate. First, we show concertedness is an outcome, not a mechanism. Second, we demonstrate that selection regime and structural interdependencies are critical to the outcome of competing mechanisms. Third, we argue that both developmentally coupled and uncoupled brain architectures can both be adaptive, but we contest the assumption that developmental coupling is necessarily a persistent evolutionary constraint. Finally, our model provides a way to integrate patterns of brain evolution, life history and environmental heterogeneity. We of course acknowledge that our model is a simplification of a highly complex biological phenomenon. While the assumptions we make in constructing the model are intended to help us clearly explore previous verbal arguments, several extensions are possible. These include the addition of heterogeneous distributions of energetic costs, costs of functional integration to mirror energetic costs of long-range axon, and a total brain size ceiling imposed by allometric scaling with body size, which is itself under selection. Constraints imposed by co-evolution between brain and body size are of particular interest, as they may limit energetic investments in brain tissue, for example, increasing the strength of trade-offs between components. This could alter the outcome of competition between *D* values. Nevertheless, we hope the general approach taken can trigger greater formalisation of evolutionary hypotheses in this field, and work that further refines our computational models.

## Acknowledgements

We thank Corina Logan, for instigating this collaboration, and the Castiglione family for facilitating discussions.

## Author contributions

SA, AC and SHM devised the model, interpreted the output and wrote the manuscript. SA produced the code to implement the model.

