## Supplementary Figures for "An agent-based model clarifies the importance of functional and developmental integration in shaping brain evolution"

### List of supplementary figures

**Figure S1:** A simplified, pictorial depiction of the model. Pages 4-5.

**Figure S2:** Evolution of 'mosaicism' under alternative conditions, full comparisons, run with 5% mutation size and 100 generations [sister to Figure 1]. Page 6.

**Figure S3:** Evolution of 'mosaicism' under alternative conditions, full comparisons, run with 50% mutation size and 100 generations [extension of Figure 1]. Page 7.

**Figure S4:** Evolution of 'mosaicism' under alternative conditions, full comparisons, run with 50% mutation size and 10 generations [extension of Figure 1]. Page 8.

**Figure S5:** Generation number at convergence during simulations of competition between evolving populations with different *D* values under alternative conditions [companion to Figure 2]. Page 9.

**Figure S6:** Competition between evolving populations with different *D* values under alternative conditions, full comparisons, run with 5% mutation size and 100 generations [sister to Figure 2]. Page 10.

**Figure S7:** Competition between evolving populations with different *D* values under alternative conditions, full comparisons, run with 50% mutation size and 100 generations [extension of Figures 2 and 3]. Page 11.

**Figure S8:** Competition between evolving populations with different *D* values under alternative conditions, full comparisons, run with 50% mutation size and 10 generations [extension of Figures 2 and 3]. Page 12.

**Figure S9:** Competition between evolving populations with different *D* values under tailored environmental conditions, run with 5% mutation size, showing the average size of each brain component and the frequency of competing *D* values [extension of Figures 2 and 3]. Page 13.

**Figure S10:** Competition between evolving populations with different *D* values under tailored environmental conditions, full comparisons, run with 5% mutation size and 100 generations [extension of Figures 2 and 3]. Page 14.

**Figure S11:** Competition between evolving populations with different *D* values under tailored environmental conditions, full comparisons, run with 50% mutation size and 100 generations [extension of Figures 2 and 3]. Page 15.

**Figure S12:** Competition between evolving populations with different *D* values under tailored environmental conditions, full comparisons, run with 50% mutation size and 10 generations [extension of Figures 2 and 3]. Page 16.

**Figure S13:** Conditions at convergence of simulations of competition between evolving populations with different *D* values in a randomly varying environment, under different life history conditions [companion to Figure 2]. Page 17.

**Figure S14:** Competition between evolving populations with different *D* values in a varying environment, under different life history conditions, full comparisons, run with 5% mutation size and 100 generations [sister to Figure 4]. Page 18.

**Figure S15:** Competition between evolving populations with different *D* values in a varying environment, under different life history conditions, full comparisons, run with 50% mutation size and 100 generations [extension of Figure 4]. Page 19.

**Figure S16:** Competition between evolving populations with different *D* values in a varying environment, under different life history conditions, full comparisons, run with 50% mutation size and 10 generations [extension of Figure 4]. Page 20.

**Figure S17:** Selected, representative, individual simulations showing fluctuations in population frequencies over generations for a 5% mutation size (A-D) or a 50% mutation size (E-H) [extension of Figure 4]. Page 21.

**Figure S18:** Subtle effects of the size of environmental fluctuations on competition between evolving populations with different *D* values in a varying environment [extension of Figure S16]. Page 22.

**Figure S19:** Relationship between population frequencies between partially mosaic brains (*D*=0.5) and fully mosaic (*D*=0), or concerted brains (*D*=1) from simulations in varying environmental conditions and a 5% mutation size (A) or 50% mutation size (B) [extension of Figure 4]. Page 23.

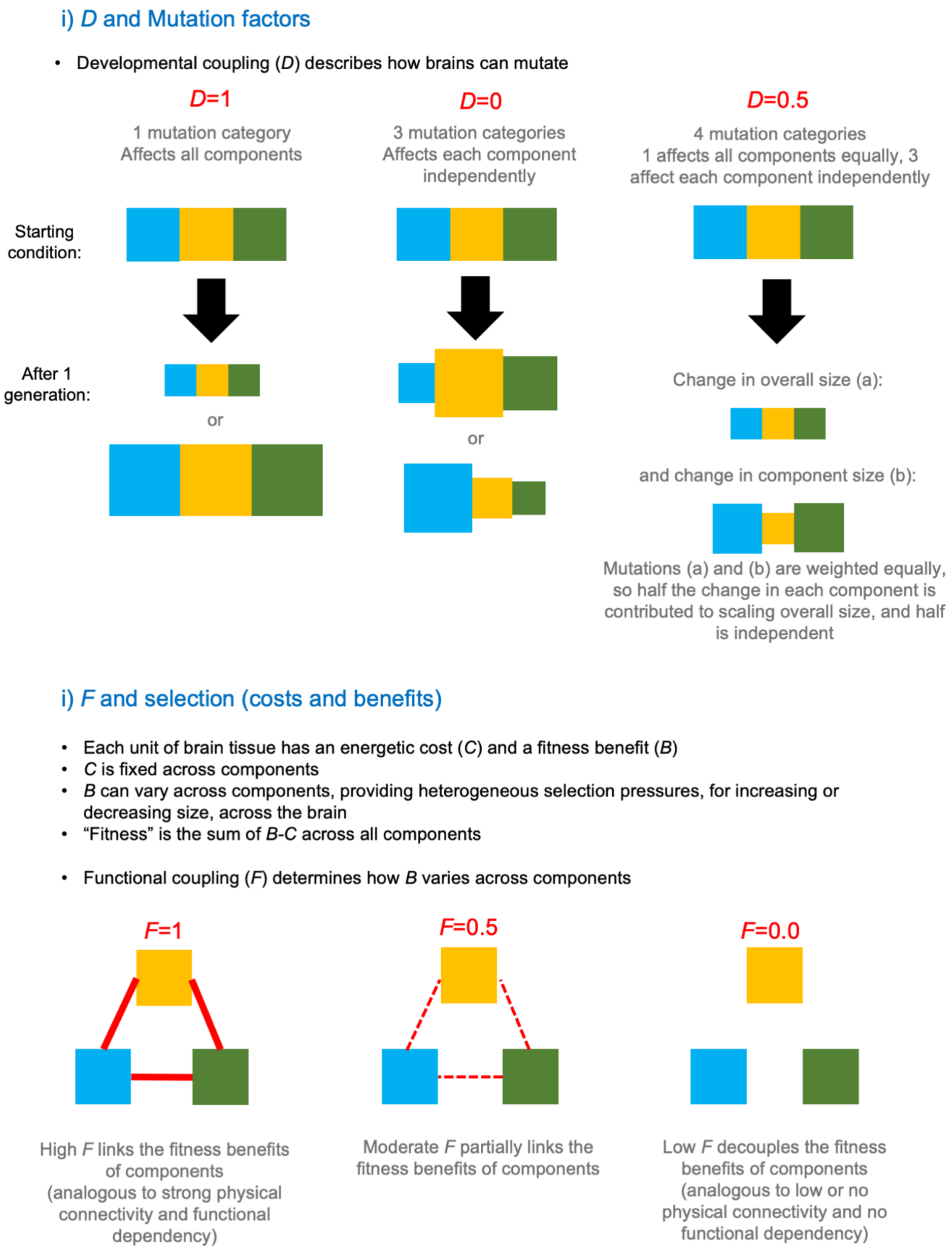


**Figure S1:** A simplified, pictorial depiction of the model.

#### iii) Life history of a strategy

- Each individual lives for a set number of generations
- Each individual breeds once a set number of generations, producing a set number of offspring
- Generations can overlap

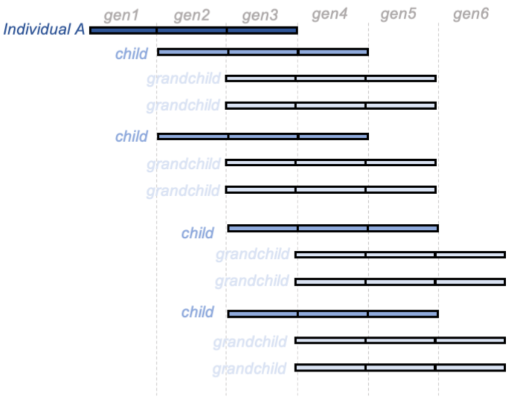


*e.g.* a population derived from Individual A living for 3 generations, with each individual producing 2 offspring per generation after 1 generation to reach maturity. Great-grandchildren not shown for clarity.

- Population size is kept constant, meaning excess individuals are culled each generation, see (iv) for details.
- This creates 'competition' between strategies with selection determined by the environment imposed by the distribution of *B* across components

#### iv) Selection and population turnover

- Each individual produces a set number of offspring each of which are 'mutated', as described in (i)
- Variation in overall brain size, and the size of each brain component creates variation in 'fitness' (see ii)
- At each generation, all the individuals are ranked by their 'fitness'

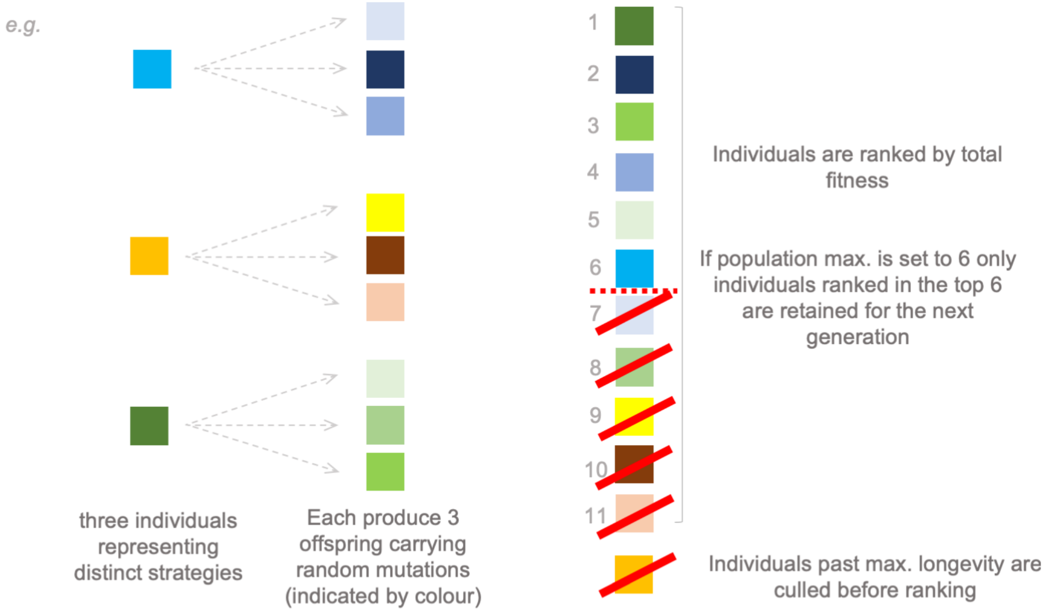


- This process is repeated across a set number of generations. From a starting point of equal representation of strategies, the change in the frequency of a strategy indicates its selective advantage.

**Figure S1 cont'd:** A simplified, pictorial depiction of the model.

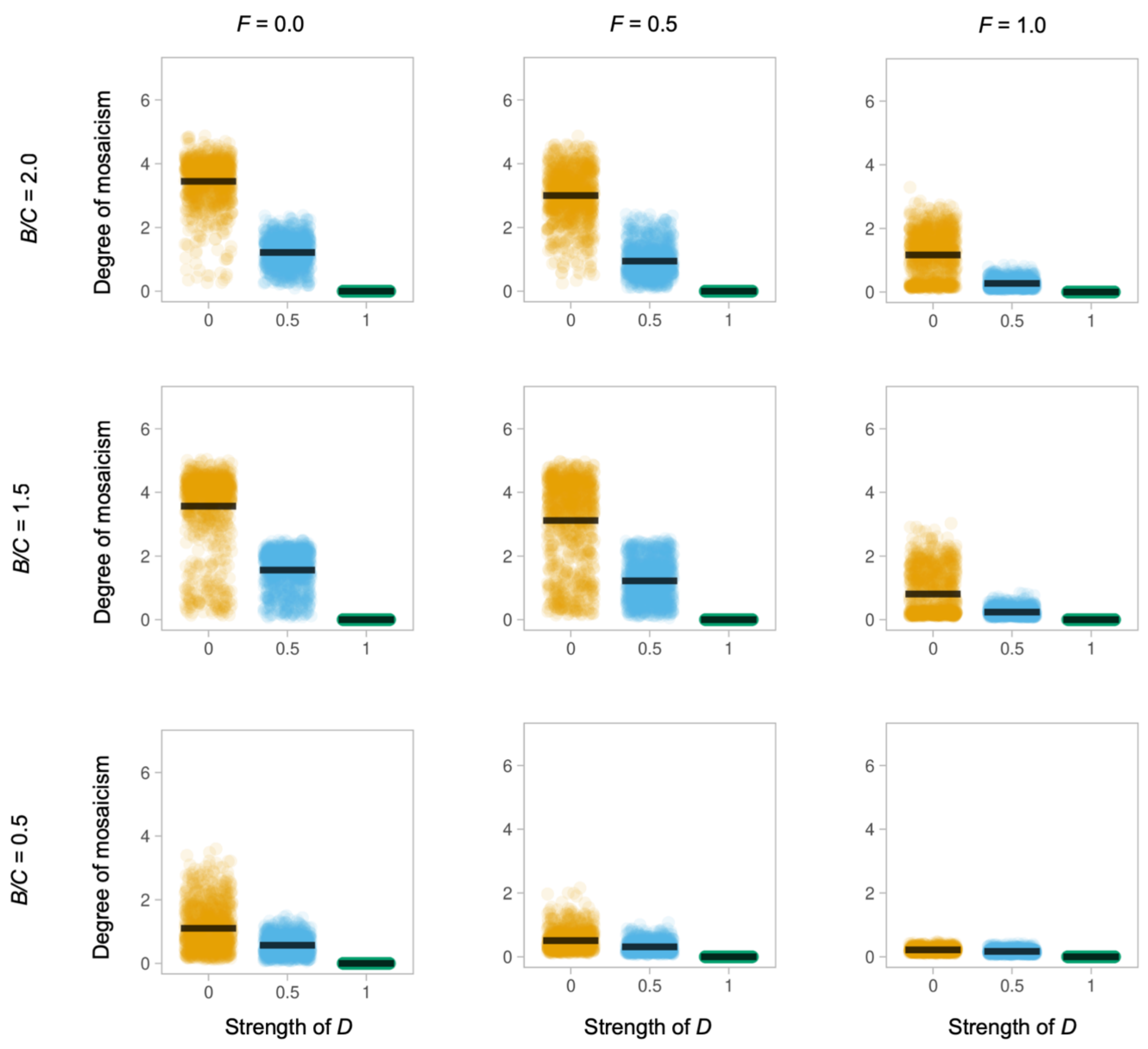


**Figure S2:** Evolution of 'mosaicism' under alternative conditions, with a mutation step size of 5%. Each plot depicts the 'degree of mosaicism' (y-axis, defined as the natural log of the ratio between the largest brain component and the smallest brain component in each individual, averaged across the population) as a function of developmental coupling *D* (x-axis) at the end of a 100-generation simulation run, compiled over 1,000 simulation runs, for a population of 100 individuals with an identical *D*, under different environments, defined by their functional coupling *F*, and their average benefit to cost ratio [sister to Figure 1].

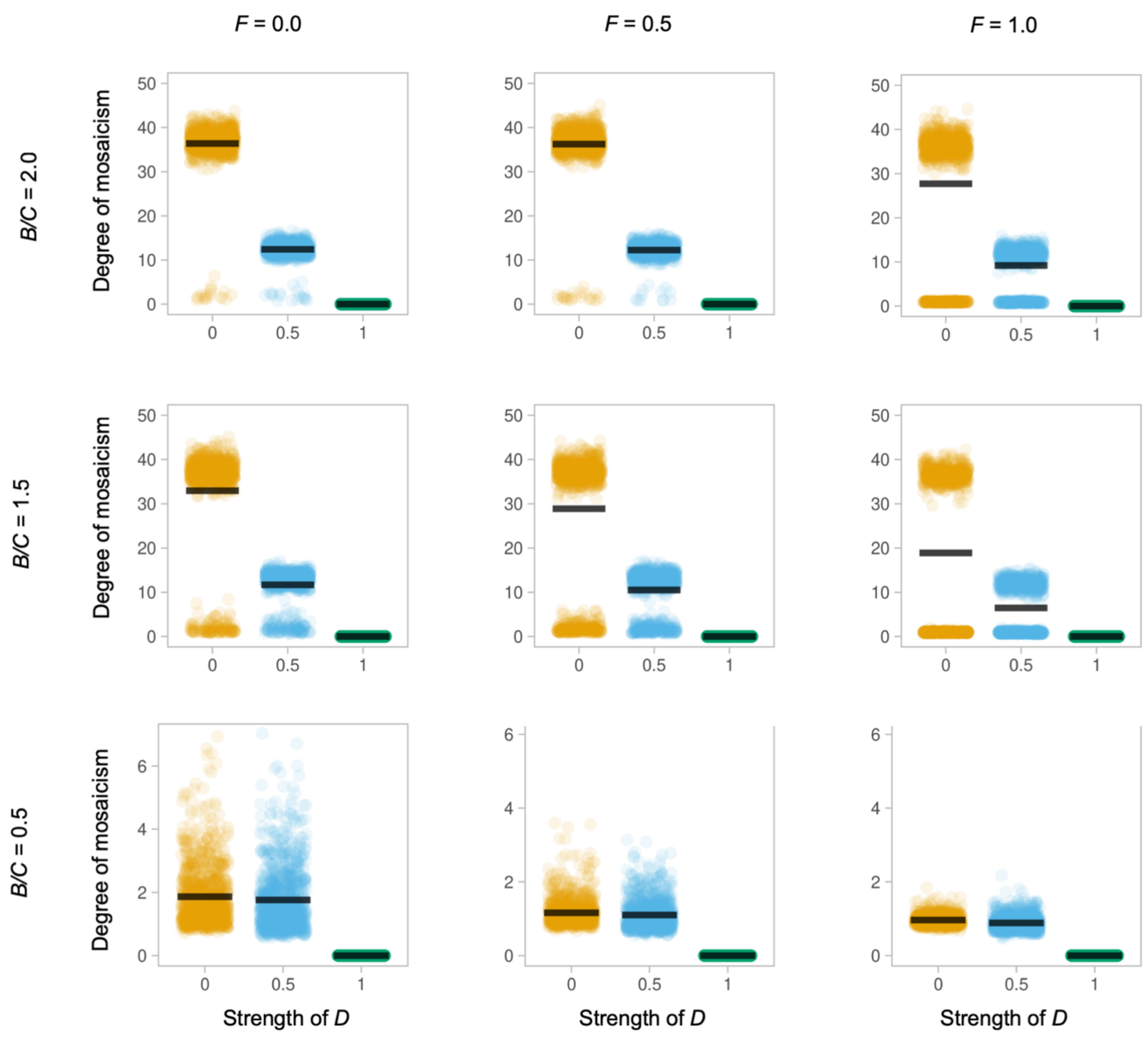


**Figure S3:** Evolution of 'mosaicism' under alternative conditions, with a mutation step size of 50%. Each plot depicts the 'degree of mosaicism' (y-axis, defined as the natural log of the ratio between the largest brain component and the smallest brain component in each individual, averaged across the population) as a function of developmental coupling *D* (x-axis) at the end of a 100-generation simulation run, compiled over 1,000 simulation runs, for a population of 100 individuals with an identical *D*, under different environments, defined by their functional coupling *F*, and their average benefit to cost ratio. Interestingly our model predicts a step change in levels of mosaicism under some scenarios (e.g. row 2), which can be attributed to the mutational effect size (compare with row 2 in Figure S2). This implies large effect mutations could allow rapid 'jumps' in mosaicism. [extension of Figure 1].

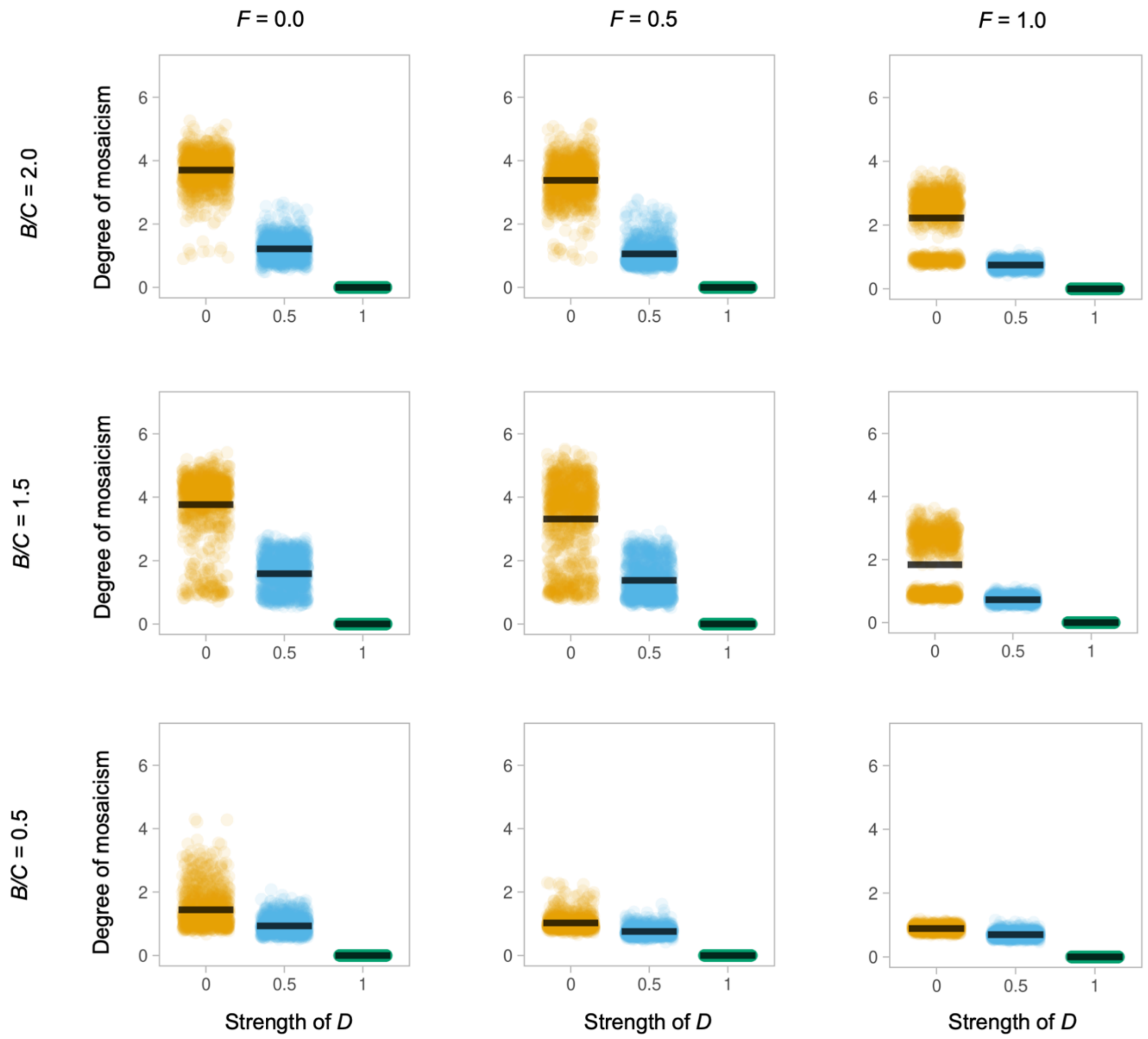


**Figure S4:** Evolution of 'mosaicism' under alternative conditions, with a mutation step size of 50%. Each plot depicts the 'degree of mosaicism' (y-axis, defined as the natural log of the ratio between the largest brain component and the smallest brain component in each individual, averaged across the population) as a function of developmental coupling *D* (x-axis) at the end of a 10-generation simulation run, compiled over 1,000 simulation runs, for a population of 100 individuals with an identical *D*, under different environments, defined by their functional coupling *F*, and their average benefit to cost ratio [extension of Figure 1].

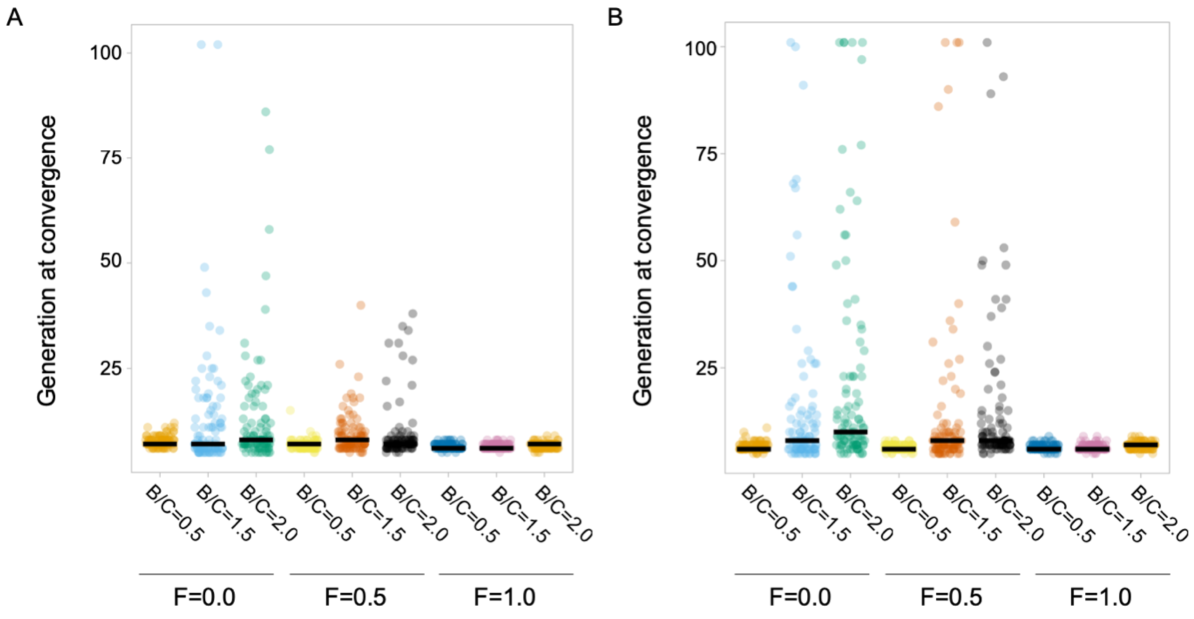


**Figure S5:** Generation number at convergence during simulations of competition between evolving populations with different *D* values under alternative *F* and *B/C* conditions, and a 5% (A) or 50% (B) mutation step size. Convergence is defined as the generation at which one population represents 100% of individuals in the environment. Iterations at generation 150 had not yet converged on a single *D* value. Convergence is slower when $F<1$ and $B>1$, where the difference in population success is less pronounced, and so multiple populations persist for longer. Convergence is also slower with larger mutation sizes, as population means vary more between generations. [companion to Figure 2].

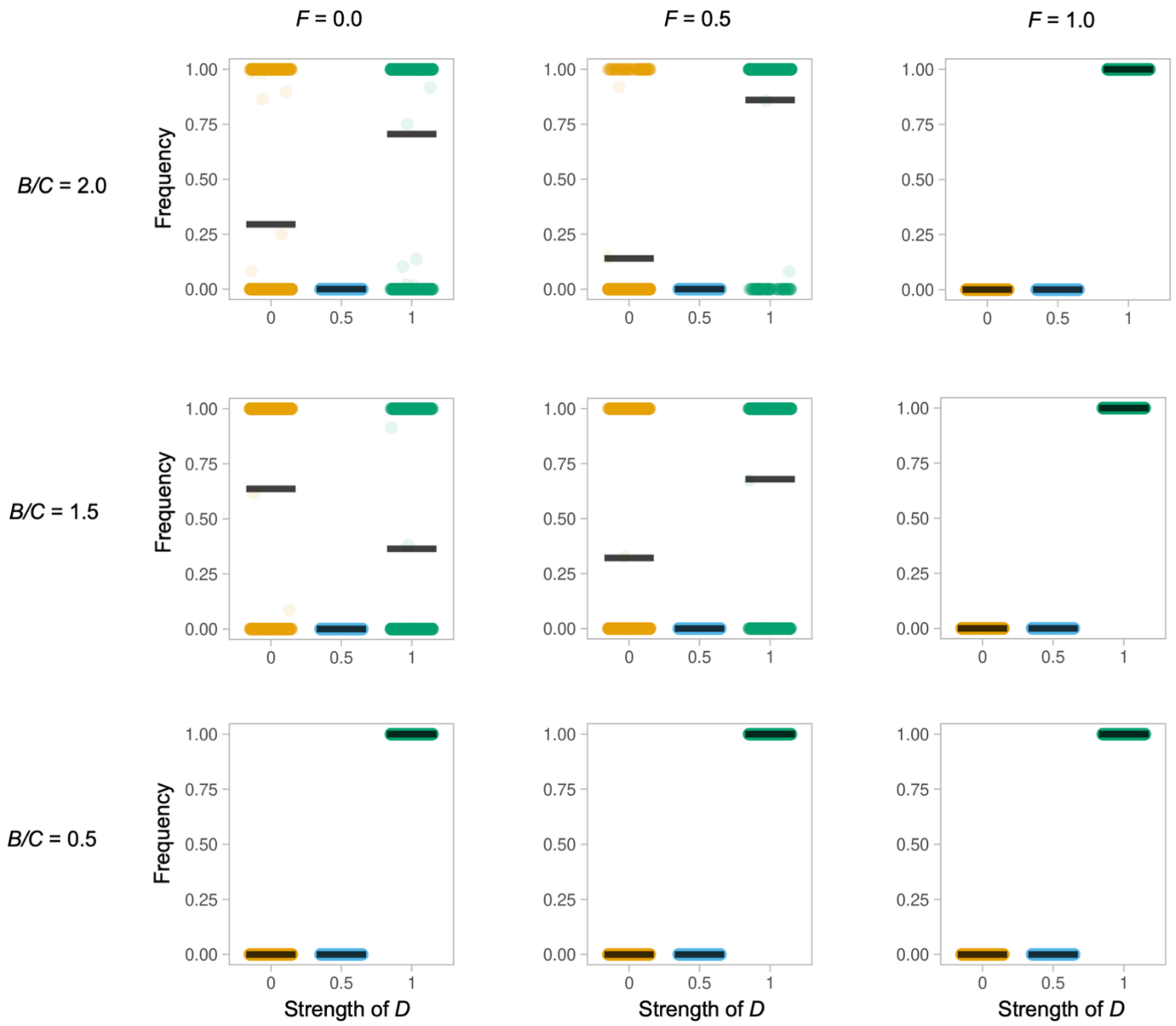


**Figure S6:** Competition between evolving populations with different *D* values under alternative conditions, and a 5% mutation step size. Each plot depicts the ratio of that population to the total population (y-axis) as a function of developmental coupling *D* (x-axis) at the end of a 100-generation simulation run, compiled over 1,000 simulation runs, under alternative environments defined by their functional coupling *F,* and their average benefit to cost ratio. Populations are initialised such that there are 100 fully mosaic individuals (*D*=0.0), 100 hybrid individuals (*D*=0.5) and 100 concerted individuals (*D*=1.0). [sister to Figure 2].

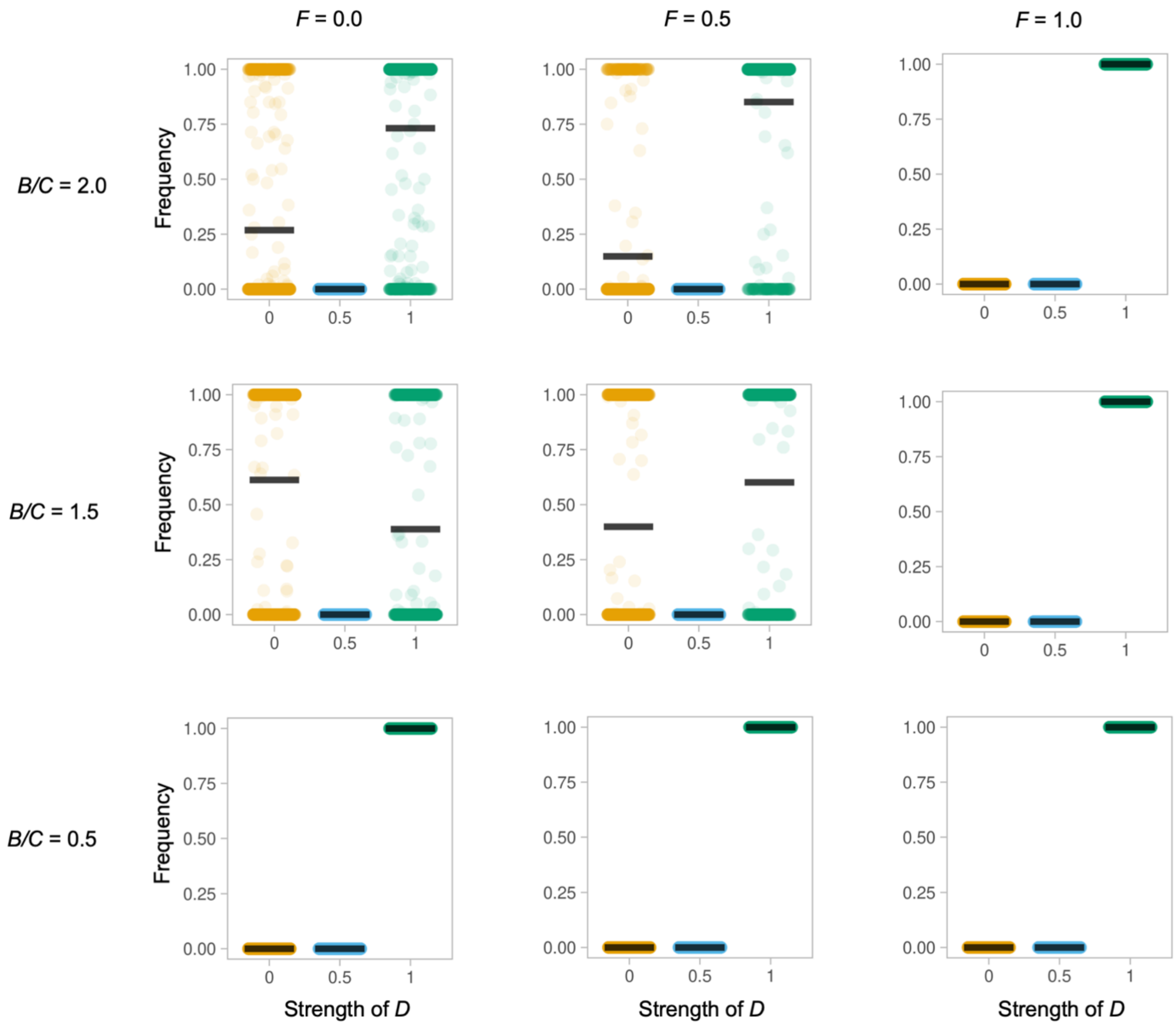


**Figure S7:** Competition between evolving populations with different *D* values under alternative conditions, and a 50% mutation step size. Each plot depicts the ratio of that population to the total population (y-axis) as a function of developmental coupling *D* (x-axis) at the end of a 100-generation simulation run, compiled over 1,000 simulation runs, under alternative environments defined by their functional coupling *F,* and their average benefit to cost ratio. Populations are initialised such that there are 100 fully mosaic individuals (*D*=0.0), 100 hybrid individuals (*D*=0.5) and 100 concerted individuals (*D*=1.0) [extension of Figures 2 and 3].

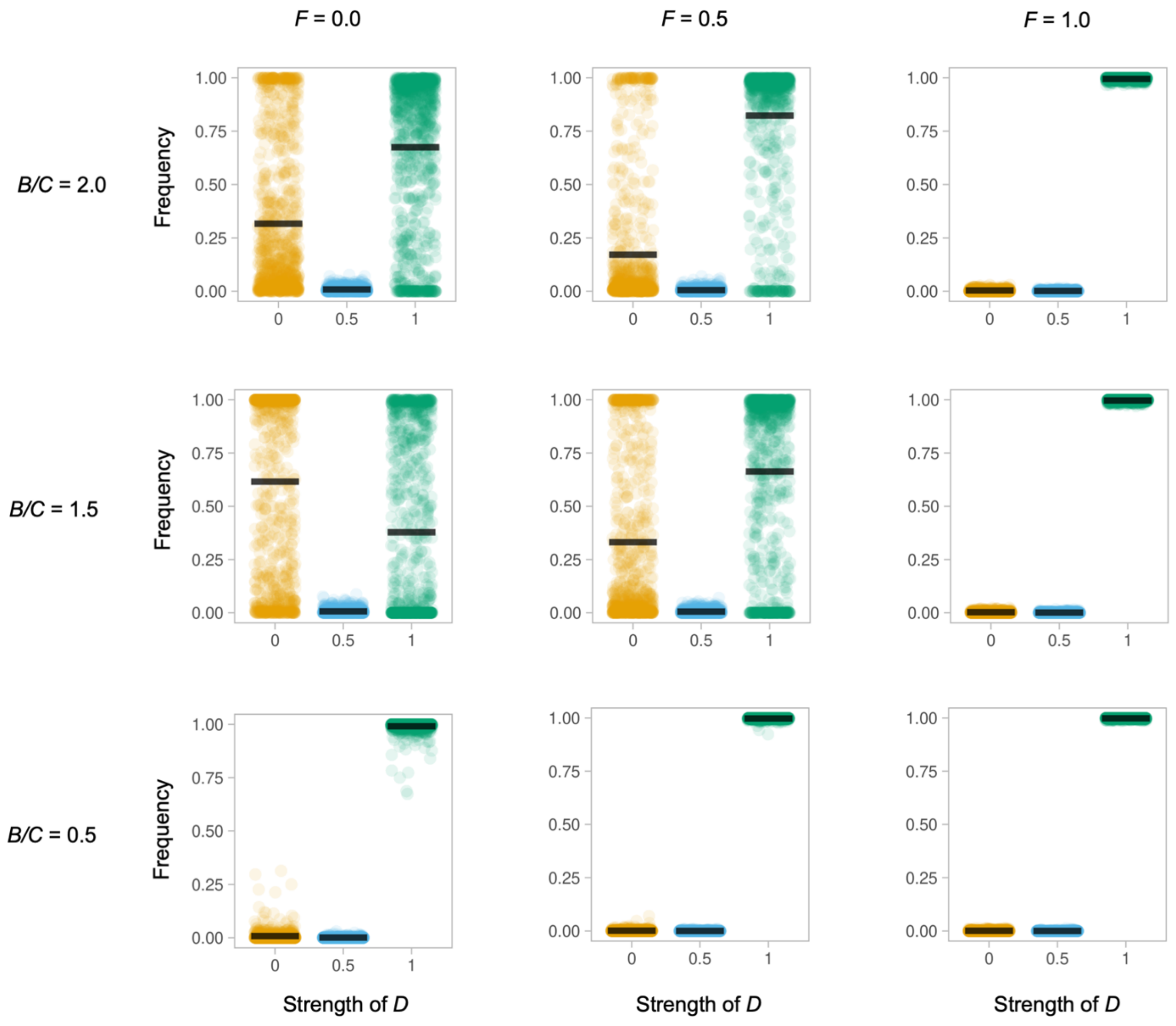


**Figure S8:** Competition between evolving populations with different *D* values under alternative conditions, and a 50% mutation step size. Each plot depicts the ratio of that population to the total population (y-axis) as a function of developmental coupling *D* (x-axis) at the end of a 10-generation simulation run, compiled over 1,000 simulation runs, under alternative environments defined by their functional coupling *F,* and their average benefit to cost ratio. Populations are initialised such that there are 100 fully mosaic individuals (*D*=0.0), 100 hybrid individuals (*D*=0.5) and 100 concerted individuals (*D*=1.0) [extension of Figures 2 and 3].

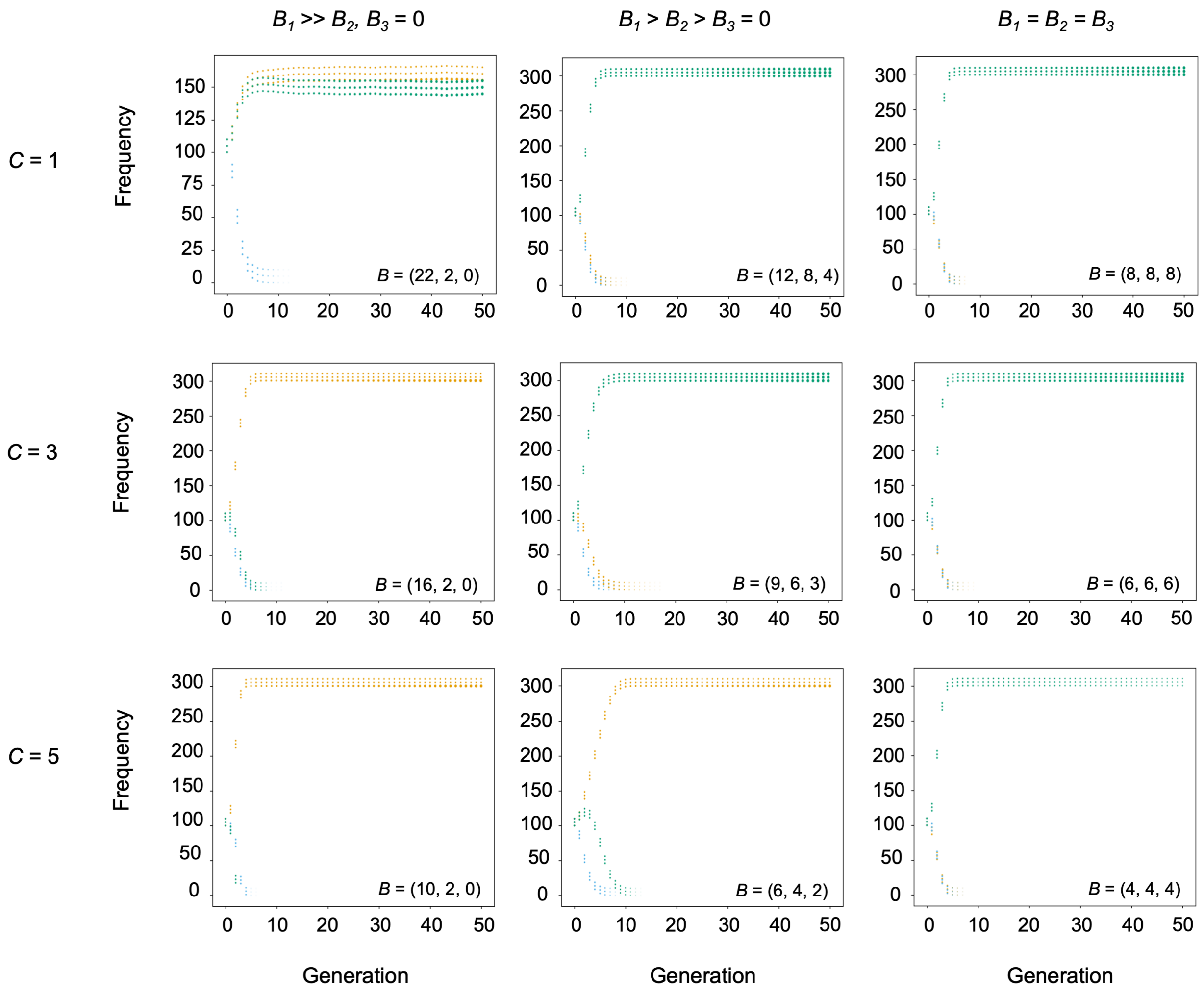


**Figure S9:** Competition between evolving populations with different *D* values under alternative conditions, with a mutation step size of 5%. Each plot depicts the frequency of individuals within a population (y-axis) as a function of developmental coupling *D* (colour coded: *D*=0 in gold, *D*=0.5 in blue, *D*=1.0 in green) during the first 50 generations of simulation runs, compiled over 1,000 simulation, with "hand-crafted" environments depicted in the figure to explore specific situations of interest. The plots show the average size of each component in the 3-component brain across the population of each *D* value [extension of Figure 4].

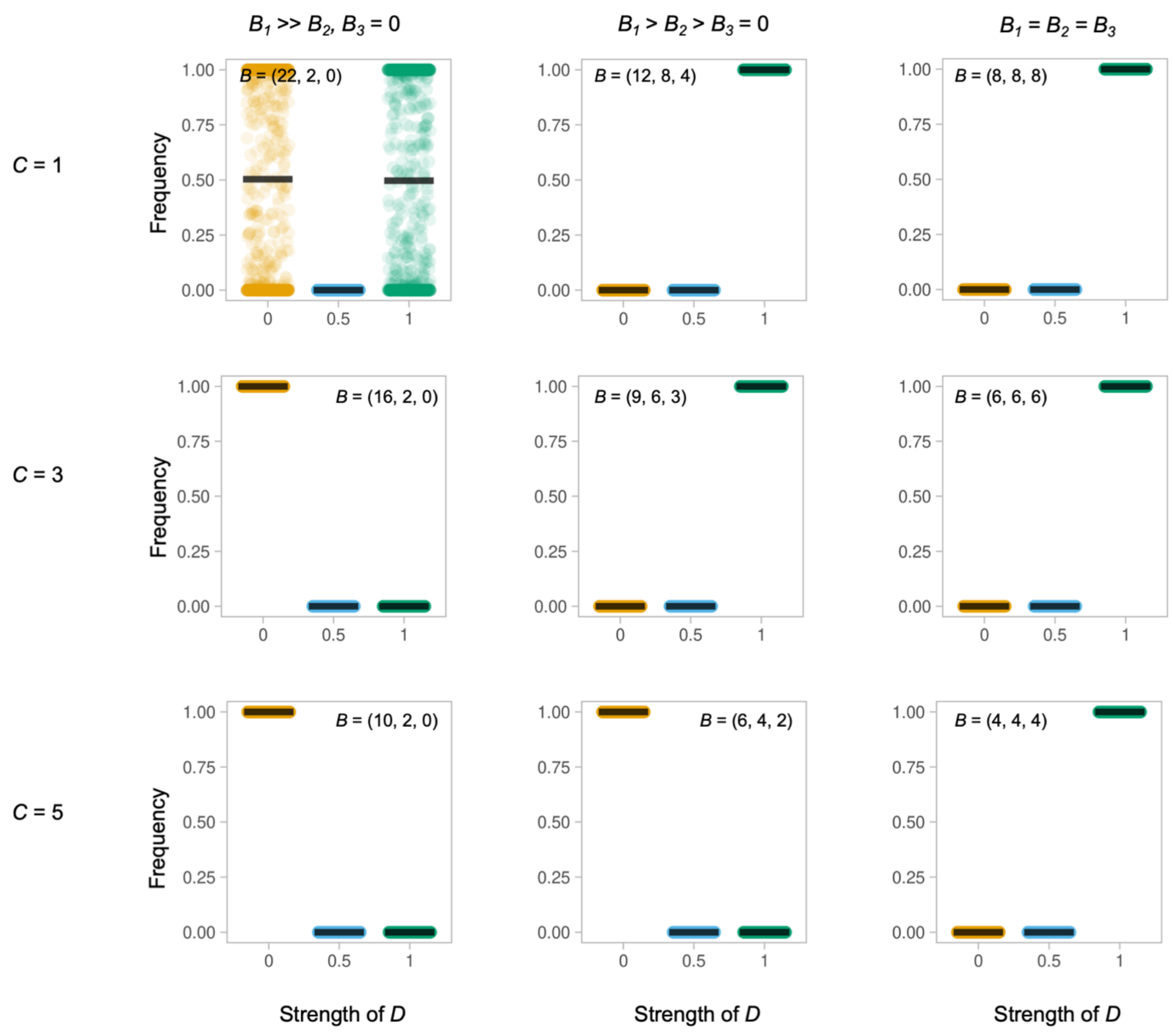


**Figure S10:** Competition between evolving populations with different $D$ values under alternative conditions, with a mutation step size of 5%. Each plot depicts the ratio of that population to the total population (y-axis) as a function of developmental coupling $D$ (x-axis) at the end of a 100-generation simulation run, compiled over 1,000 simulation runs, with "hand-crafted" environments depicted in the figure to explore specific situations of interest [extension of Figures 2 and 3].

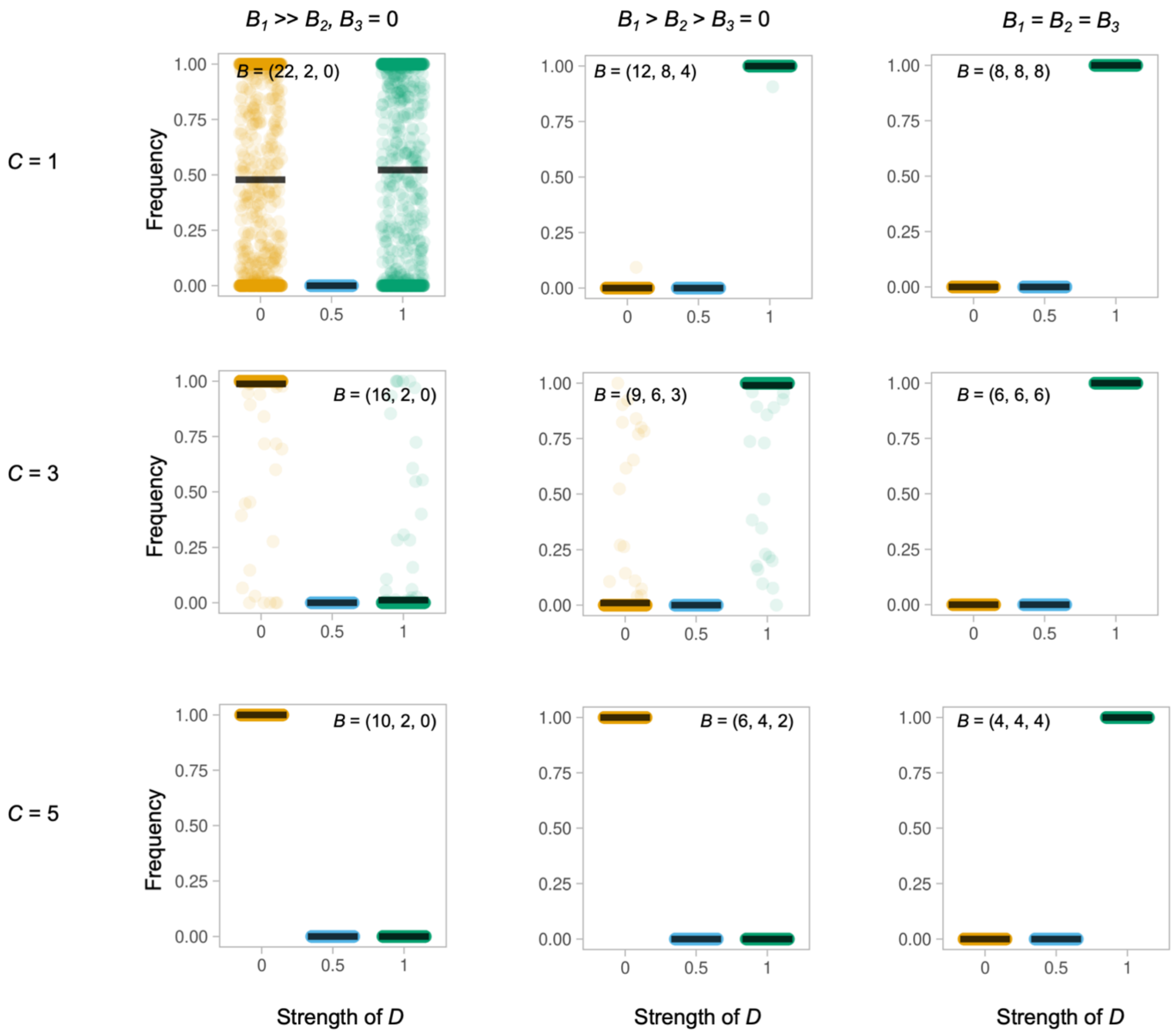


**Figure S11:** Competition between evolving populations with different *D* values under alternative conditions, with a mutation step size of 50%. Each plot depicts the ratio of that population to the total population (y-axis) as a function of developmental coupling *D* (x-axis) at the end of a 100-generation simulation run, compiled over 1,000 simulation runs, with "hand-crafted" environments depicted in the figure to explore specific situations of interest [extension of Figures 2 and 3].

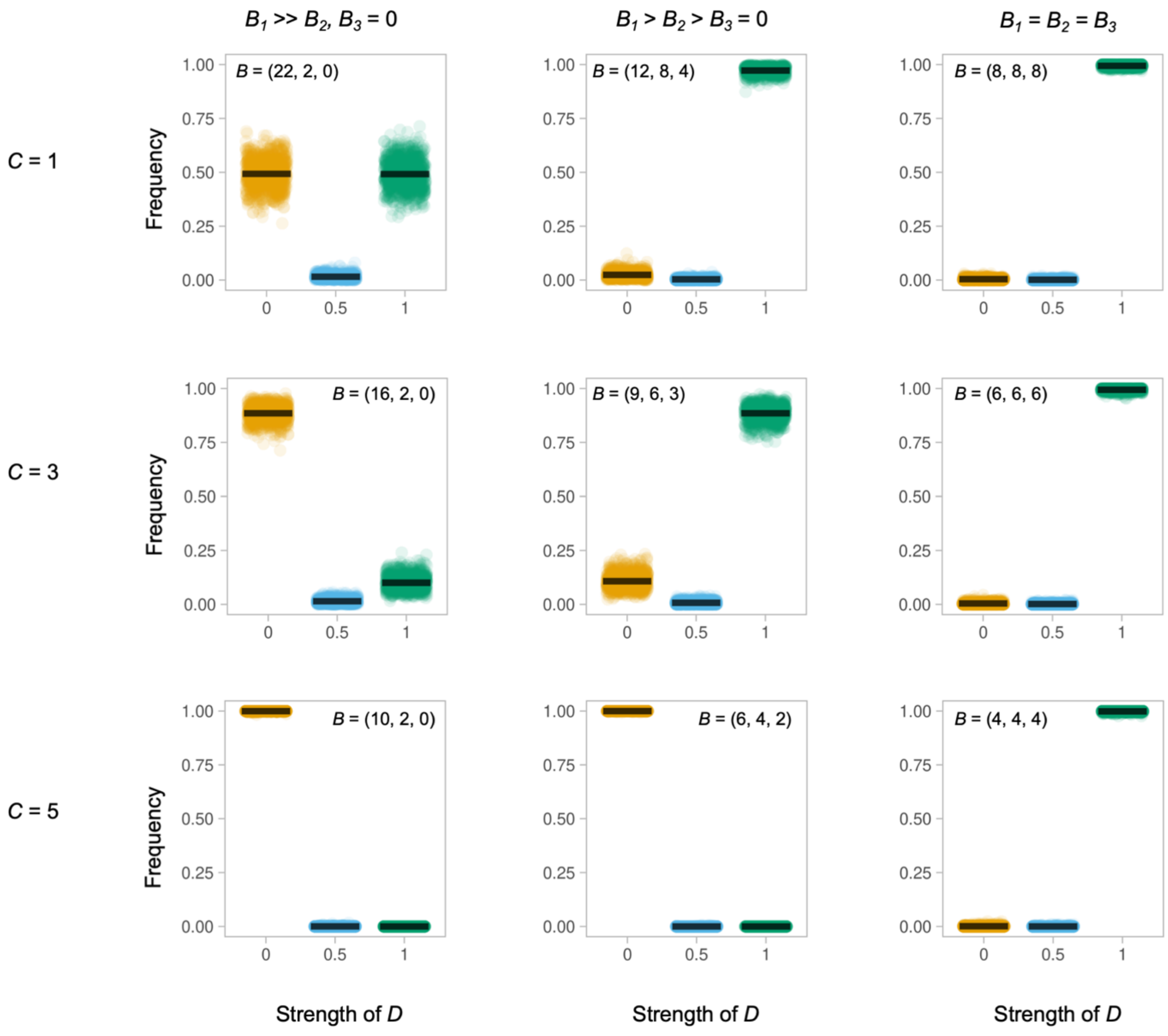


**Figure S12:** Competition between evolving populations with different *D* values under alternative conditions, with a mutation step size of 50%. Each plot depicts the ratio of that population to the total population (y-axis) as a function of developmental coupling *D* (x-axis) at the end of a 10-generation simulation run, compiled over 1,000 simulation runs, with "hand-crafted" environments depicted in the figure to explore specific situations of interest [extension of Figures 2 and 3].

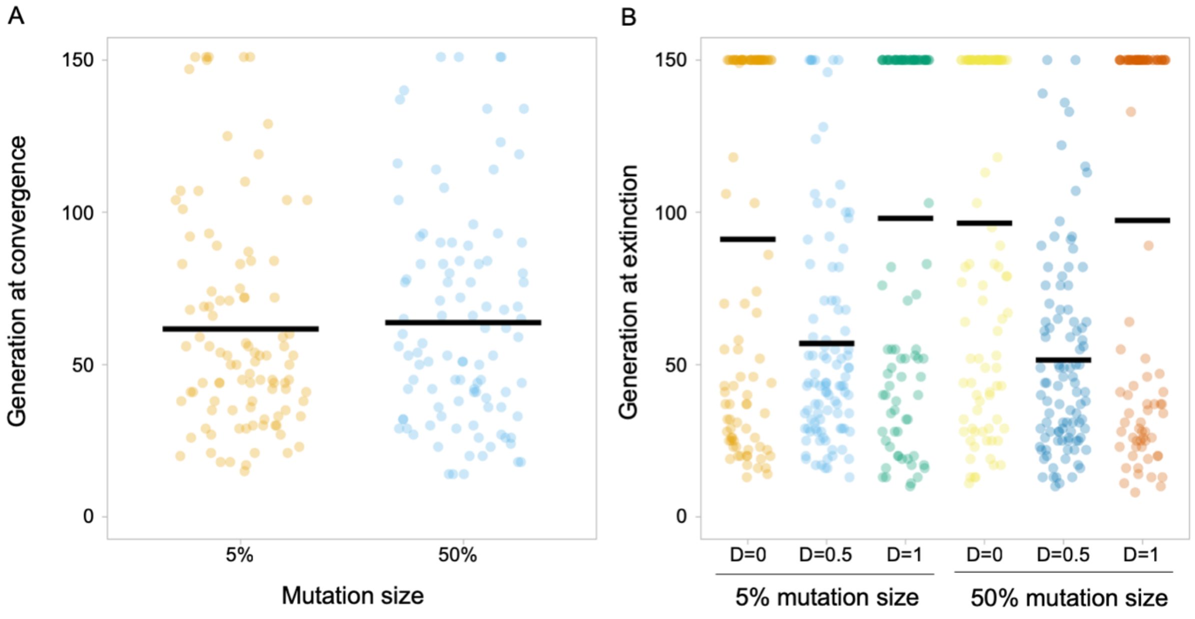


**Figure S13:** Conditions at convergence of simulations of competition between evolving populations with different *D* values in a randomly varying environment, under different life history conditions. Convergence is defined as the generation at which one population represents 100% of individuals in the environment, extinction is defined as the generation at which a population drops to zero individuals. Iterations at generation 100 had not yet converged on a single population [companion to Figure 2].

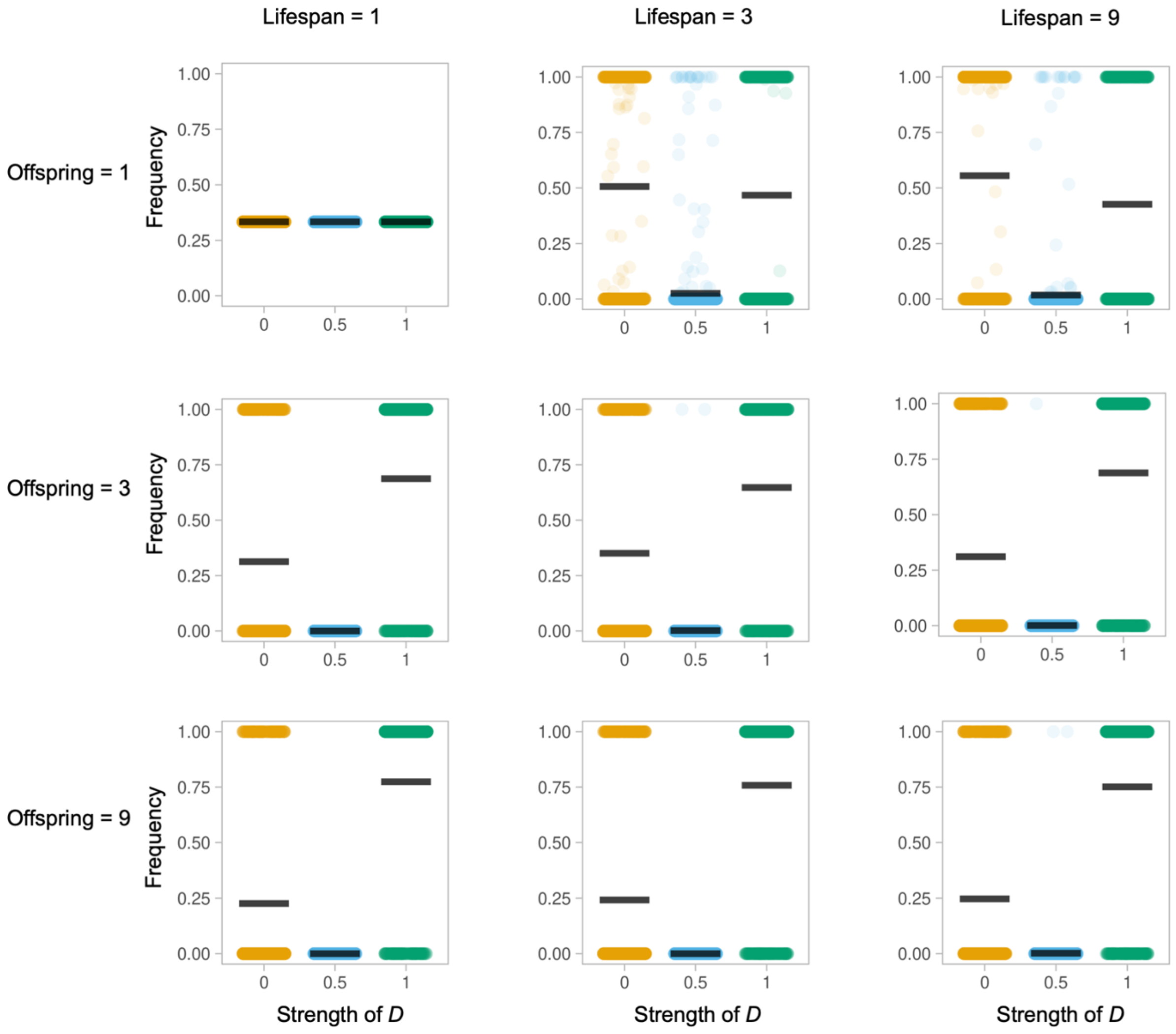


**Figure S14:** Competition between evolving populations with different *D* values in a varying environment, under different life history conditions (lifespan, or number of generations an individual persists, and offspring number, which are generated every generation), with a 5% mutation step size. Each plot depicts the ratio of that population to the total population (y-axis) as a function of developmental coupling *D* (x-axis) at the end of a 150-generation simulation run, compiled over 1,000 simulation runs; every 2 generations the environment is replaced, using a uniform random distribution for cost [0.5, 5], max component benefit [1.0, 10], and *F* functional coupling [0, 1]. Populations are initialised such that there are 100 fully mosaic evolvers (*D*=0.0), 100 hybrid evolvers (*D*=0.5) and 100 concerted evolvers (*D*=1.0) [sister to Figure 5].

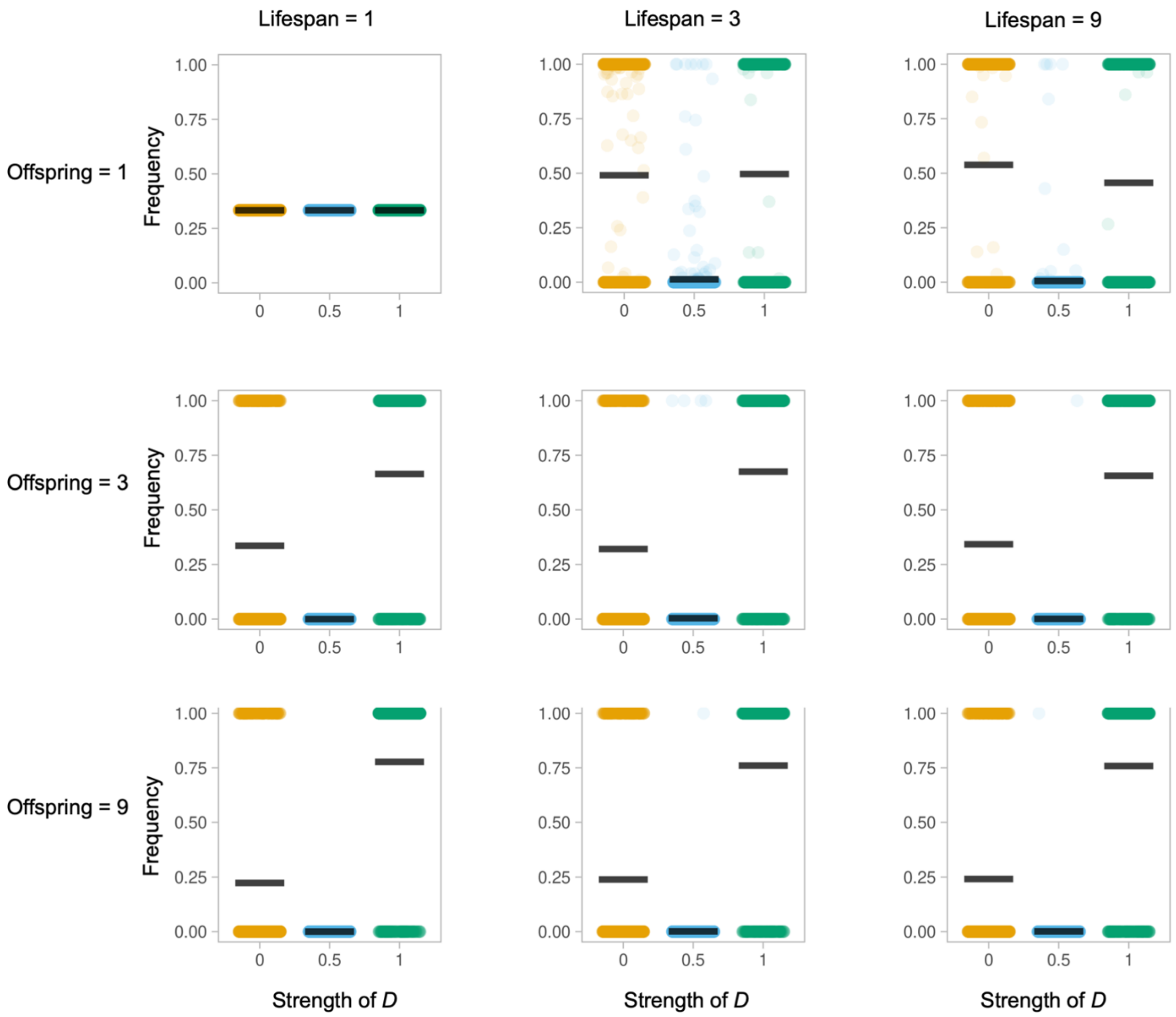


**Figure S15:** Competition between evolving populations with different *D* values in a varying environment, under different life history conditions (lifespan, or number of generations an individual persists, and offspring number, which are generated every generation), with a 50% mutation step size. Each plot depicts the ratio of that population to the total population (y-axis) as a function of developmental coupling *D* (x-axis) at the end of a 150-generation simulation run, compiled over 1,000 simulation runs; every 2 generations the environment is replaced, using a uniform random distribution for cost [0.5, 5], max component benefit [1.0, 10], and *F* functional coupling [0, 1]. Populations are initialised such that there are 100 fully mosaic evolvers (*D*=0.0), 100 hybrid evolvers (*D*=0.5) and 100 concerted evolvers (*D*=1.0) [sister to Figure 5].

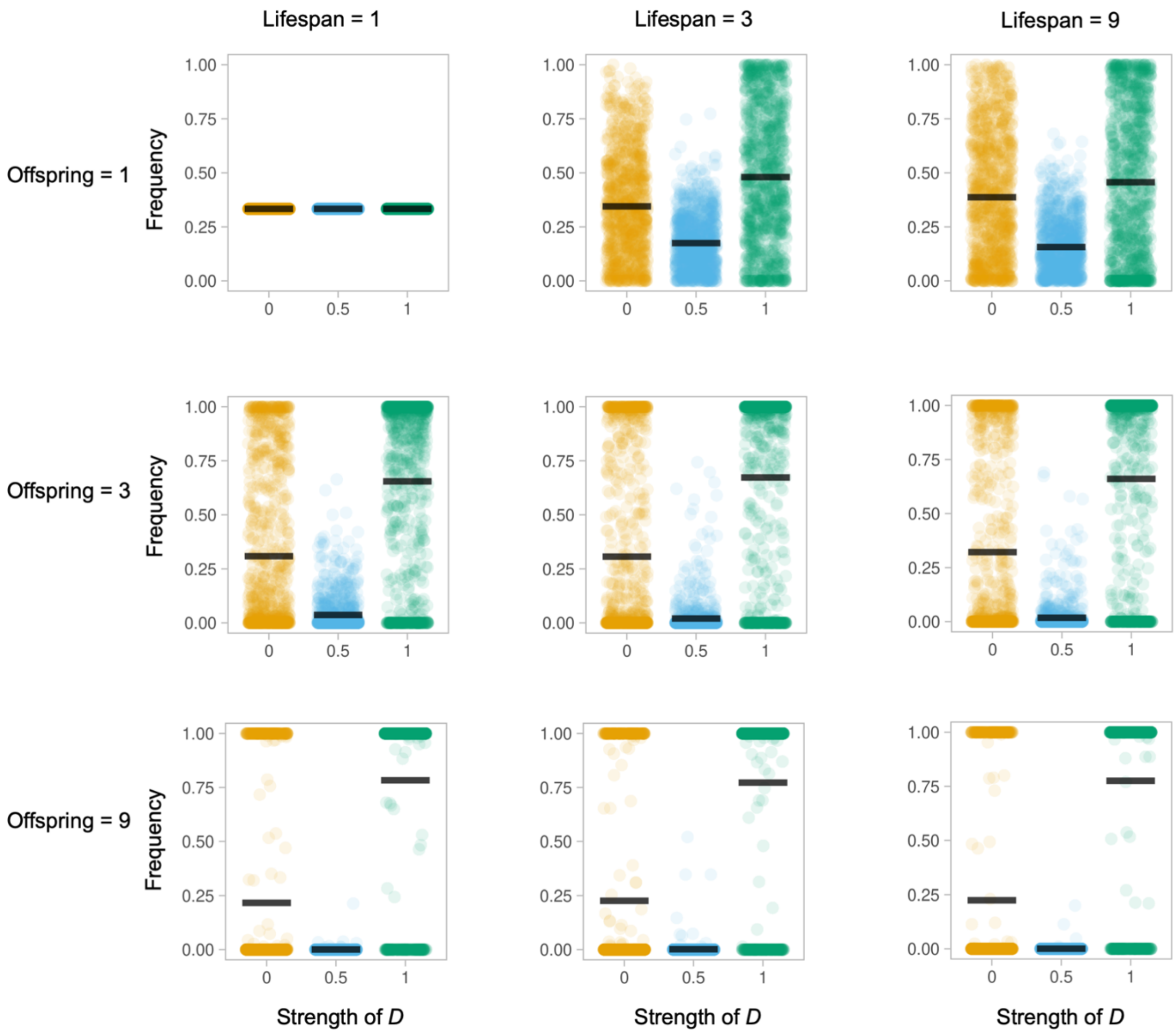


**Figure S16:** Competition between evolving populations with different *D* values in a varying environment, under different life history conditions (lifespan, or number of generations an individual persists, and offspring number, which are generated every generation), with a 5% mutation step size. Each plot depicts the ratio of that population to the total population (y-axis) as a function of developmental coupling *D* (x-axis) at the end of a 10-generation simulation run, compiled over 1,000 simulation runs; every 2 generations the environment is replaced, using a uniform random distribution for cost [0.5, 5], max component benefit [1.0, 10], and *F* functional coupling [0, 1]. Populations are initialised such that there are 100 fully mosaic evolvers (*D*=0.0), 100 hybrid evolvers (*D*=0.5) and 100 concerted evolvers (*D*=1.0) [sister to Figure 5].

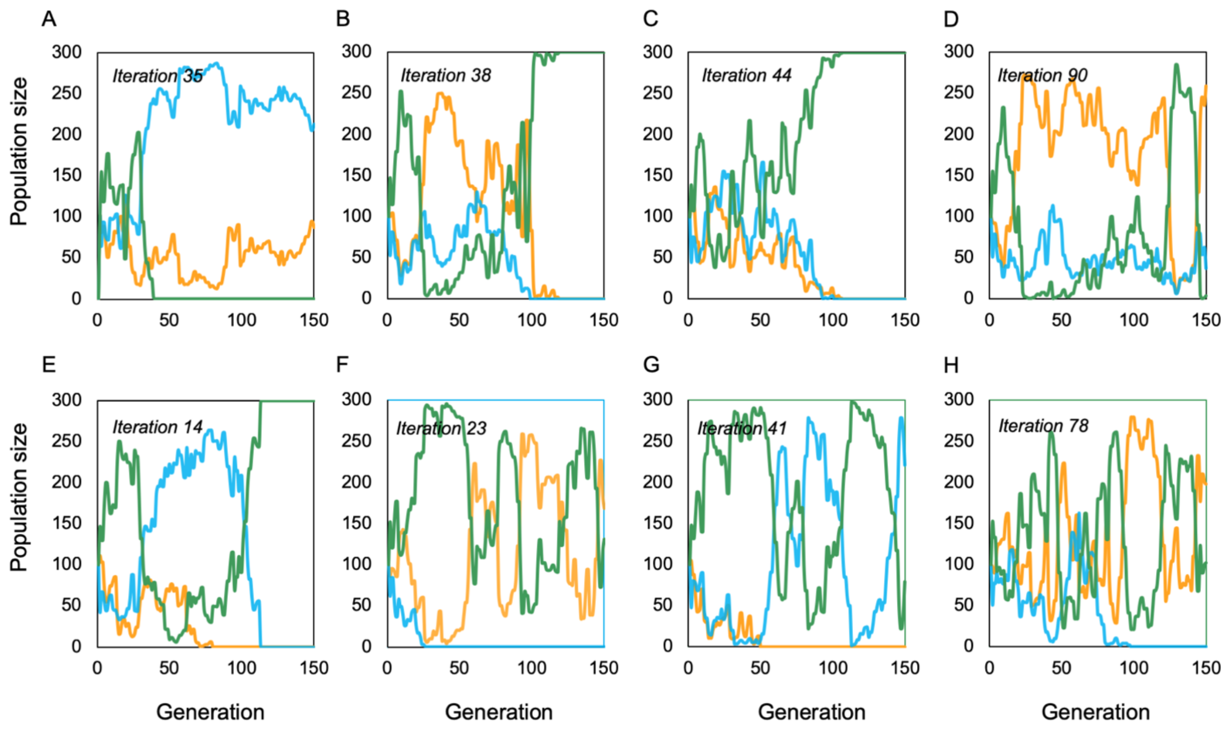


**Figure S17:** Selected, representative, individual simulations showing fluctuations in population frequencies over generations for a 5% mutation size (A-D) or a 50% mutation size (E-H). Colours indicate the *D* value where yellow is *D* =0, where blue is *D* =0.5, and where green is *D* =1. These examples illustrate the central conclusion from this set of simulations, that multiple population can persist across time together. We interpret this as suggesting allelic variation for concerted and mosaic evolution could be maintained in a population under fluctuating environmental conditions, and hence be available to selection. [extension of Figure 5].

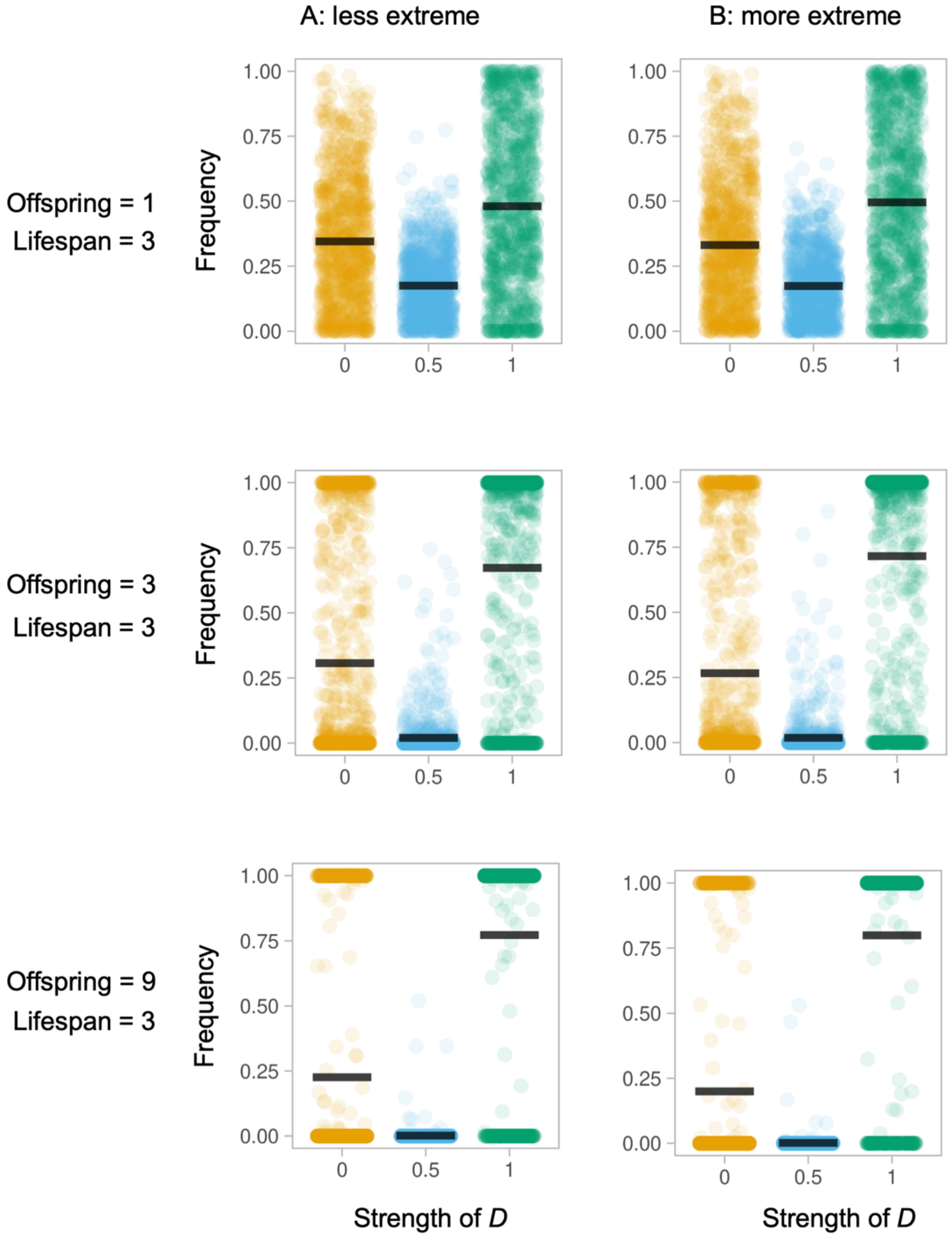


**Figure S18:** Competition between evolving populations with different *D* values in a varying environment, under selected life history conditions, as described for Figure S15 (column A), but with an increased distribution for cost [0, 10] and max average benefit [0, 30] creating more 'extreme' environmental fluctuations (column B). During early generations of the simulations, more extreme environmental fluctuations increase the probability of a concerted population being successful, as indicated by a significant interaction between *D* and the size fluctuation size category ($t = 8.986$, $p<0.001$), but with a low effect size. [extension of Figure S16].

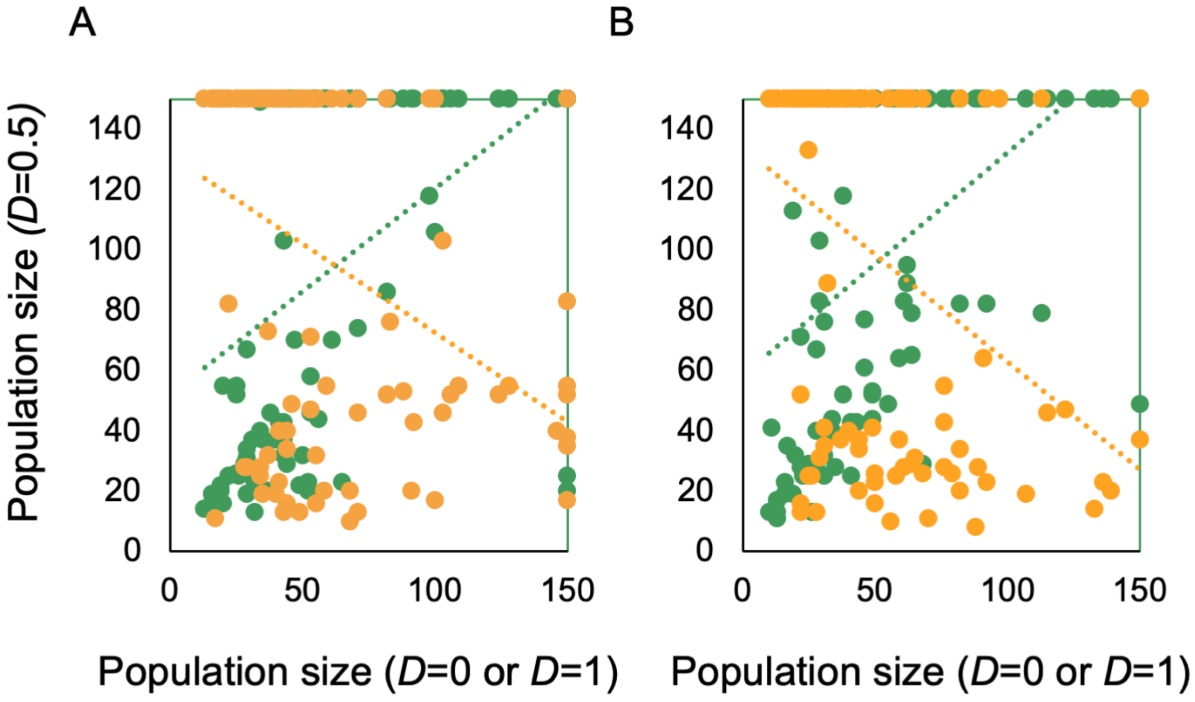


**Figure S19:** Relationship between population frequencies between partially mosaic brains (*D*=0.5) and fully mosaic (*D*=0), or concerted brains (*D*=1) from simulations in varying environmental conditions and a 5% mutation size (A) or 50% mutation size (B). The plot shows that *D*=0 and *D*=0.5 are more commonly favoured in the same environments (shown in green) than *D*=0 and *D*=1.0 (shown in yellow) [extension of Figure 5].
